# Macrophage Inflammasome Activation Drives Anti-inflammatory Responses via Neutrophil-Derived ASC Speck Transfer and Gasdermin-Dependent Caspase-1 Inhibition

**DOI:** 10.1101/2025.01.22.634381

**Authors:** Annamaria Pedoto, Juan M. Lozano Gil, Mai E. Nguyen-Chi, Georges Lutfalla, Diana García-Moreno, María Luisa Cayuela, Sylwia D. Tyrkalska, Victoriano Mulero

## Abstract

Inflammasomes are critical components of the innate immune system, traditionally associated with pro-inflammatory responses. While the inflammasome in macrophages has been extensively studied and linked to pyroptosis and cytokine release, the neutrophil inflammasome remains poorly understood. Neutrophil inflammasome activation drives unique outcomes such as NETosis and robust IL-1β production without inducing pyroptosis. In contrast, macrophage inflammasome activation promotes pyroptosis and the release of cytokines like IL-1α and IL-18. Here, using zebrafish models of Salmonella enterica serovar Typhimurium infection and COVID-19-associated cytokine storm syndrome (CSS), we reveal opposing roles of neutrophil and macrophage inflammasomes: neutrophil inflammasomes are pro-inflammatory, whereas macrophage inflammasomes mediate anti-inflammatory responses. We show that Gasdermin E- and Ninjurin-1-mediated cell death in neutrophils releases ASC specks, which are taken up by macrophages. This transfer negatively regulates caspase-1 activity via the C-terminal domain of Gasdermin E, promoting the resolution of inflammation. To our knowledge, this is the first study to reveal the anti-inflammatory role of the macrophage inflammasome, document the *in vivo* transfer of ASC specks between neutrophils and macrophages, and uncover the negative feedback regulation of the inflammasome by the C-terminal domain of Gsdme. These results offer novel insights into potential therapeutic approaches for hyperinflammatory diseases like bacterial infections and COVID-19-associated CSS.

## INTRODUCTION

The inflammasomes are multiprotein complexes situated in the cytosol of the immune cells, mostly macrophages, neutrophils and dendritic cells, that are crucially involved for the activation of the inflammatory response and cell death (*1*). They typically consist of a sensor protein, the adaptor protein apoptosis-associated speck-like protein containing a caspase-recruitment domain (ASC), and the proinflammatory caspase, caspase-1 (*2, 3*). Assembly of the inflammasome is a well-organized process comprising an inflammatory signaling cascade. It can be triggered by recognizing of a variety of stimuli that are associated with infection or cellular stress (*4*). Inflammasome assembly requires homotypic CARD–CARD or PYD–PYD interactions between its components inducing oligomerization (*5, 6*). When the ligand is being detected, the sensor is relieved from an inhibitory state and recruits ASC to form a multimeric complex called speck. The sensor protein can include NLRs, AIM2 or pyrin depending on the type of danger signal involved in the process and they are categorized based on their structural characteristics (*7*). Then, ASC recruits procaspase-1 (proCASP1) into the complex, which next is converted into active CASP1 that cleaves pro-IL-1β and pro-IL-18 to active cytokines and activate pore-forming Gasdermin D (GSDMD) to induce a form of a programmed cell death called pyroptosis (*8, 9*). Moreover, it has been shown that activation of the inflammasome can lead to the generation of an eicosanoid storm through the induction of cytosolic phospholipase A2 (PLA2) in resident peritoneal macrophages or neutrophils (*10, 11*).

Neutrophils and macrophages are key innate immune cells that form the first line of defense against pathogens (*12, 13*). Neutrophils, which constitute ∼70% of circulating leukocytes, are rapidly recruited to inflammation sites, where they eliminate microbes through phagocytosis, degranulation, and the release of neutrophil extracellular traps (NETs) (*14–16*). In contrast, macrophages reside in tissues, where they clear debris and regulate inflammation (*17*). These cells can polarize into two main subsets: pro-inflammatory M1 macrophages and anti-inflammatory M2 macrophages, with their balance being critical for resolving inflammation and tissue repair (*18*).

Although both neutrophils and macrophages express functional inflammasomes, the neutrophil inflammasome remains largely understudied. Historically, inflammasome activation and IL-1β production were primarily attributed to macrophages and dendritic cells, with neutrophils considered passive recipients of pro-inflammatory cytokines (*16*). However, recent studies have revealed that neutrophils can activate NLRP3, NLRC4, and NLRP12 inflammasomes during bacterial infections (*19, 20*). However, the inflammasome activation in macrophages and neutrophils leads to distinct outcomes, highlighting their functional differences. In macrophages, activation of the inflammasome causes pyroptosis, whereas in neutrophils, inflammasome activation results in GSDMD-induced formation of antimicrobial NETs and neutrophil NETosis (*16, 21, 22*). In addition, the resistance of neutrophils to pyroptotic cell death enables them to sustain IL-1β and eicosanoid production during bacterial infections (*11, 19*). This difference may be linked to the size and distribution of ASC specks: neutrophils contain multiple specks per cell, while macrophages have individual specks (*4, 23*). Additionally, in neutrophils, inflammasome activation primarily triggers IL-1β production, while in macrophages, it induces IL-1β release alongside other cytokines, such as IL-1α and IL-18 (*24*). Furthermore, while CASP1 is abundant in macrophages and drives IL-1β cleavage, in neutrophils, IL-1β cleavage is primarily mediated by serine proteases (*23*). Interestingly, neutrophils can also secrete IL-1β via an autophagy-dependent mechanism (*25*). GBPs play a central role in inflammasome activation, IL-1β production, and pyroptosis in macrophages, while they are involved only in prostaglandin biosynthesis and release in neutrophils (*11, 26*). Finally, the release of inflammasome components also differs: macrophages expel ASC specks into the extracellular space during pyroptosis (*27, 28*), while neutrophils secrete vesicles containing inflammasome components (*29*), which amplify the inflammatory response. These findings underline the distinct roles of inflammasomes in neutrophils and macrophages, emphasizing the cell-specific mechanisms that drive immune responses.

Despite the characterization of inflammasomes in both neutrophils and macrophages, all known inflammasomes to date are associated with pro-inflammatory responses, with no reported anti-inflammatory activity. Furthermore, little is known about the interactions between neutrophil and macrophage inflammasomes, especially in terms of how they may influence each other in the context of immune responses. Using the zebrafish model, here we report that inflammasomes exhibit opposing functions in these cell types: pro-inflammatory in neutrophils and anti-inflammatory in macrophages. Remarkably, specks released from neutrophils through Gasdermin E (Gsdme)- and Ninjurin-1-mediated cell death are taken up by macrophages, where they drive an anti-inflammatory response via the C-terminal Gsdme-mediated inhibition of caspase-1 activity. To our knowledge, this is the first study to reveal the anti-inflammatory role of the macrophage inflammasome, document the *in vivo* transfer of ASC specks between neutrophils and macrophages, and uncover the negative feedback regulation of the inflammasome by the C-terminal domain of Gsdme.

## RESULTS

### Inhibition of the macrophage inflammasome exacerbates inflammation in a COVID19-associated CSS model

Using a COVID-19-associated cytokine storm syndrome (CSS) model, we injected recombinant S1 domain of the SARS-CoV-2 Spike protein (S1WT) into the hindbrain ventricle (HBV) of 2 days post-fertilization (dpf) zebrafish larvae, as previously described (*30, 31*). To assess the number of *tnfa*^+^ neutrophils (N1-like) and macrophages (M1-like), we employed the reporter line *Tg(tnfa:GFP)*, crossed with either *Tg(lyz:dsRed2)* for neutrophils or *Tg(mpeg1:mCherry)* for macrophages, allowing the identification of double-positive cells. Our analysis revealed no significant differences in the number of *tnfa*^+^ neutrophils between control larvae injected with water and those injected with S1WT, both in the whole body and around the injection site (head region) (Figures 1A-1B and S1A-S1B). However, larvae injected with S1WT displayed a higher percentage of *tnfa*^+^ macrophages, both in the whole body and specifically in the head region, compared to controls (Figures 1C-1D and S1C-S1D). Despite this increase, the proportion of *tnfa*^+^ macrophages among all macrophages remained relatively low, reaching approximately 15% in the head and 10% in the whole body of S1WT-injected larvae.

**Figure 1.**
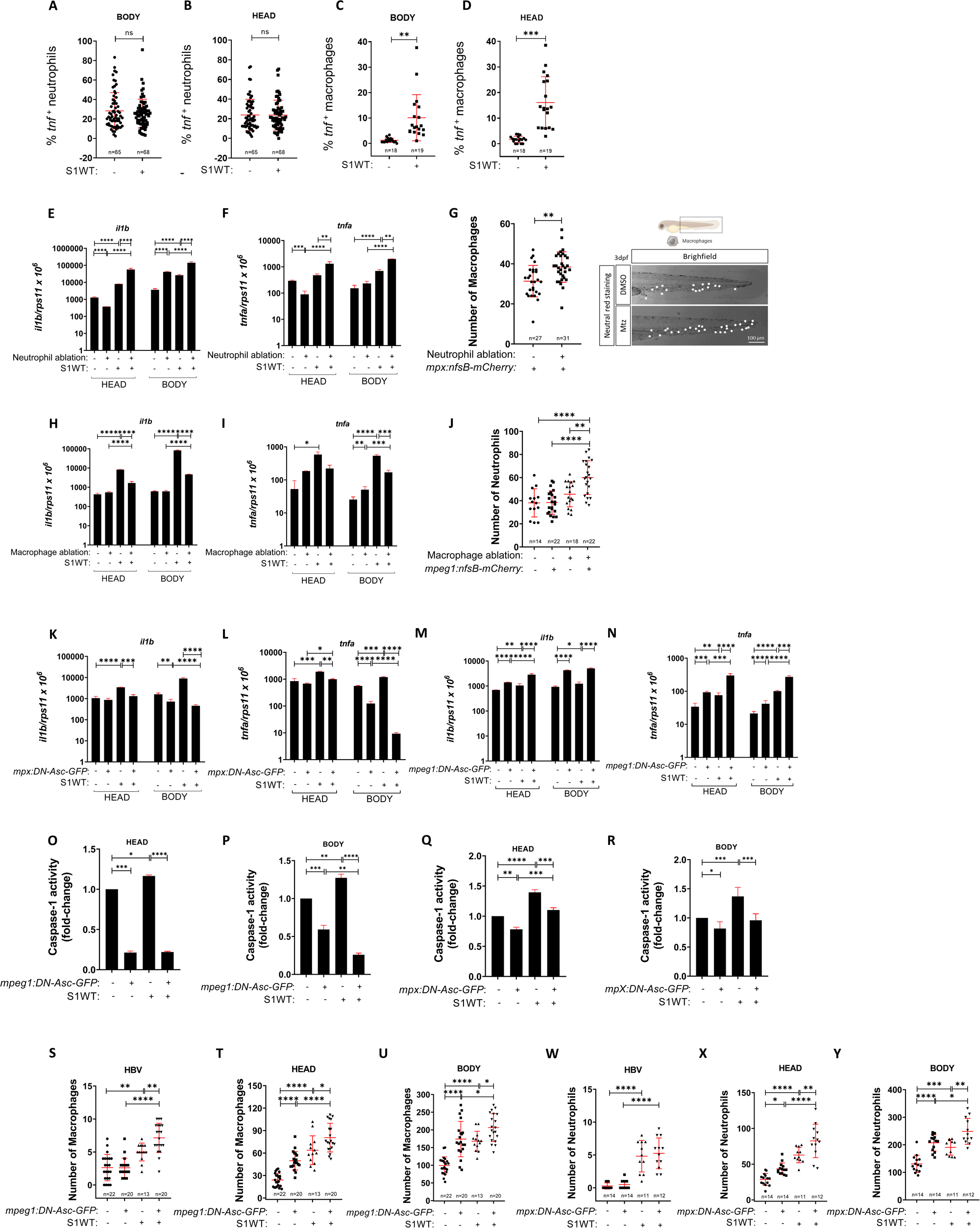
Inhibition of the macrophage inflammasome exacerbates inflammation in a COVID19-associated CSS model. Recombinant S1WT (+) or vehicle (−) were injected in the hindbrain ventricle (HBV) of 2-day postfertilization (dpf) *Tg(lyz:DsRED2)* and *Tg(tnfa:eGFP)* (A, B), Tg*(mfap4:tomato)* and *Tg(tnfa:eGFP)* (C, D), *Tg(mpx:GAL4)* and *Tg(UAS:nfsB-mCherry)* (E, F, G), Tg(mpeg1:GAL4) and Tg(UAS:nfsB-mCherry) (H, I), Tg(mpeg1:GAL4:mfap4:tomato) and Tg(UAS:nfsB-mCherry) (J), Tg(mpx:GAL4) and Tg(UAS:ascΔCARD-GFP) (K, L, Q, R), *Tg(mpeg1:GAL4)* and *Tg(UAS:DN-asc-GFP)* (M, N, O, P), *Tg(mpeg1:GAL4, mfap4:tomato)* and *Tg(UAS:DN-asc-GFP)* (S, T, U*), Tg(mpx:GAL4:lyz:DsRED2)* and *Tg(UAS:DN-asc-GFP)* (W, X, Y) larvae. The larvae were treated with 5 mM Metronidazole for specific cell ablation during 48 h (E, F, G, H, I, J). The percentage of *tnfa*^+^ neutrophils (A, B) and macrophages (C, D) were analyzed at 6 hpi using a fluorescence microscope. The transcript levels of the indicated genes (E, F, H, I, K, L, M, N) were analyzed at 12 hpi by RT-qPCR in larval head and tail. Macrophage (G, S, T, U) and neutrophil (J, W, X, Y) number and their recruitment were analyzed at 6 hpi by fluorescence microscopy or neutral red staining. Caspase-1 activity was determined at 24 hpi using a fluorogenic substrate (O, P, Q, R). Each dot represents one individual, and the means ± SEM for each group is also shown. P values were calculated using Student’s t-test or one-way analysis of variance (ANOVA) and Tukey multiple range test. *P ≤ 0.05, **P ≤ 0.01, ***P ≤ 0.001, and ****P ≤ 0.0001. auf, arbitrary units of fluorescence. Scale bars 100 µm.

We next ablated either neutrophils or macrophages, to check the role of each cell type on the inflammatory response. For that we used the GAL4-UAS system by crossing two different zebrafish transgenic lines: *Tg(uas:nfsB-mCherry)* and *Tg(mpx:gal4)* for neutrophils or *Tg(mpeg1:gal4)* for macrophages. These fish expressed bacterial nitroreductase in cell specific manner, that was able to metabolize metronidazole (MTZ) and killed the cells of interest. As we expected, S1WT-injected larvae that lacked neutrophils showed decreased levels of *il1b* and *tnfa* mRNA than the S1WT-injected control fish (Figures 1E-1F). Neutrophil ablation resulted in increased number of macrophages (Figure 1G). In sharp contrast, S1WT-injected larvae lacking macrophages showed increased levels of *il1b* and *tnfa* transcripts than S1WT-injected control fish (Figures 1H-1I). As observed before, genetic ablation of macrophages resulted in an increased number of neutrophils (Figure 1J).

To check whether these effects were mediated by cell specific inflammasome activity, we then inhibited the inflammasome in either neutrophils or macrophages by overexpressing a DN form of Asc lacking its CARD domain using GAL4-UAS system (*32*). Surprisingly, we obtained the same result seen by the depletion of neutrophils and macrophages (Figures 1K-1N), namely inhibition of the inflammasome in neutrophils led to attenuated S1WT-induced inflammation, assayed as the mRNA levels of *il1b* and *tnfa* (Figures 1K-1L). On the contrary, S1WT injected fish with inhibited inflammasome in macrophages showed increased inflammation compared with their wildtype siblings (Figures 1M-1N). We then performed caspase-1 activity assays as a readout of inflammasome activation. In both cases overexpression of DN-Asc in either neutrophils or macrophages led to decreased systemic and local caspase-1 activity in both controls and inflammation conditions (Figures 1O-1R). As expected, S1WT-injected larvae showed increased caspase-1 activity but, unexpectedly, inhibition of macrophage inflammasome resulted in much lower caspase-1 activity levels than inhibition of neutrophil inflammasome (Figure 1O-R). This result suggests that macrophages show higher inflammasome activity than neutrophils in zebrafish.

Finally, we checked the number of cells of interest upon inhibition of either macrophage or neutrophil inflammasomes. Inhibition of macrophage inflammasome resulted in increased macrophage recruitment to S1WT (Figures 1S and S1E) and monocytosis in both basal and inflammation conditions (Figures 1T-1U and S1F). In contrast, although the recruitment of neutrophils to S1WT was unaffected upon inhibition of the neutrophil inflammasome in basal and inflammatory conditions (Figure 1W and S1G), neutrophilia was apparent in larvae upon inhibition of neutrophil inflammasome in basal and inflammatory conditions (Figures 1X-1Y and S1H). These results suggest that both macrophages and neutrophils are clearer from zebrafish larvae by pyroptosis.

### Inhibition of the macrophage inflammasome exacerbates inflammation in a zebrafish infection model

To confirm the results obtained with the sterile COVID-associated CSS model, we used a bacterial infection model using *Salmonella enterica* serovar Typhimurium (ST) (*11*). The results revealed that inflammasome inhibition in either neutrophils or macrophages led to hyper susceptibility of the larvae to ST infection (Figures 2A-2B). Moreover, macrophage inflammasome inhibition resulted in hyper inflammation at 24 hpi, assayed as the transcript levels of *il1b* and *tnfa* (Figures 2C-2D). Although these results confirmed the ones obtained with the S1WT-driven inflammation, we next examined inflammation at shorter time points of the infectious process. Interestingly, at 6 hpi the inhibition of macrophage inflammasome led to lower *il1b* and *tnfa* transcript levels, whereas at 12 hpi the opposite was observed (Figures 2E-2F). Collectively, these results revealed an unappreciated role of the macrophage inflammasome in the resolution of inflammation. In sharp contrast and confirming previous results with the S1WT-driven inflammation model, neutrophil inflammasome inhibition led to decreased transcript levels of *il1b* and *tnfa* at 24 hpi (Figure 2G-H). As expected, inflammasome inhibition in both cell types decreased caspase-1 activity levels in infected larvae (Figures 2I-2J). Notably, it was found again that inhibition of macrophage inflammasome resulted in a more robust decreased of caspase-1 activity than inhibition of neutrophil inflammasome.

**Figure 2.**
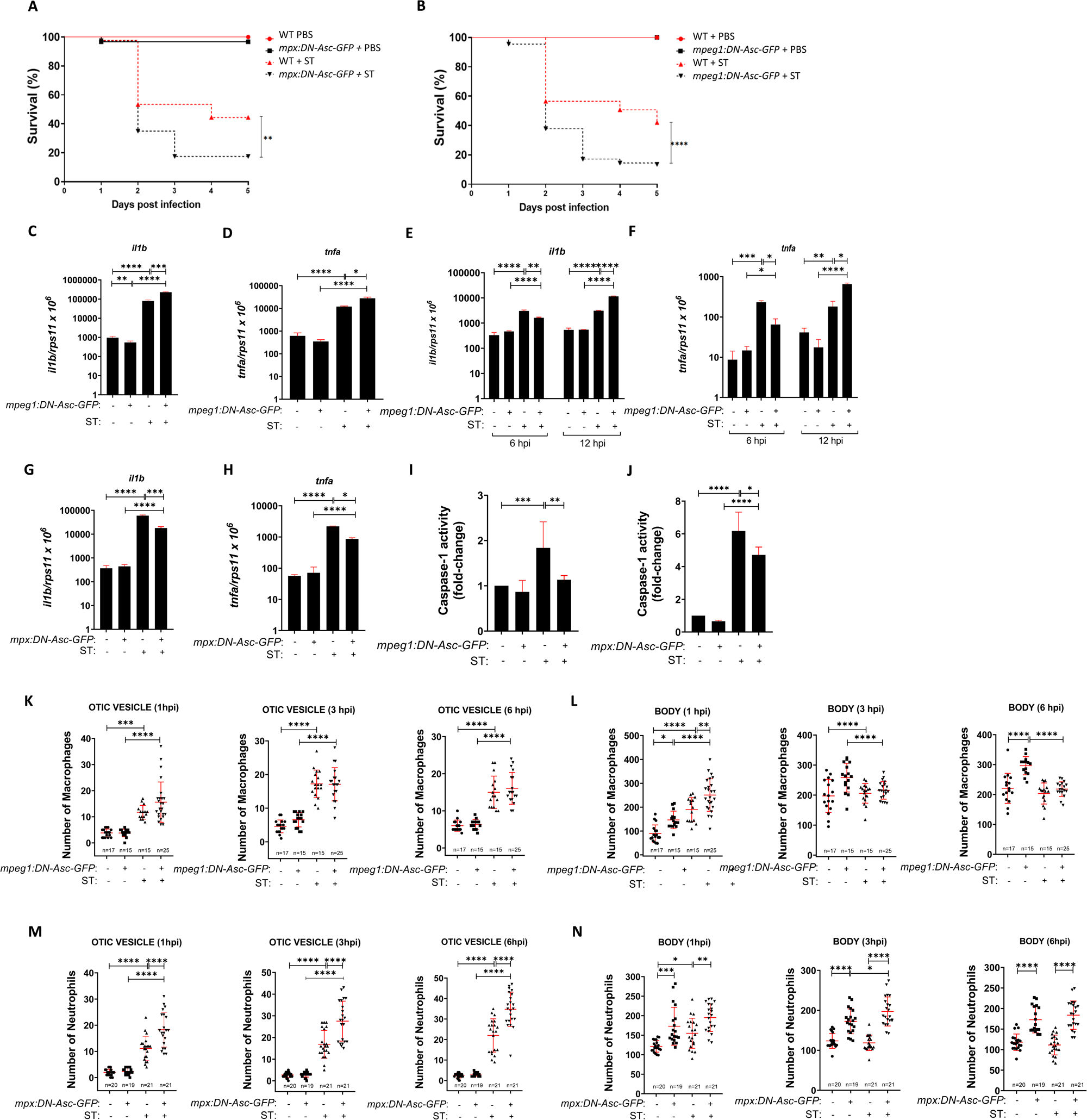
Inhibition of the macrophage inflammasome exacerbates inflammation in a zebrafish infection model. *Tg(mpx:GAL4)* and *Tg(UAS:DN-asc-GFP)* (A) and *Tg(mpeg1:GAL4)* and *Tg(UAS:DN-asc-GFP)* (B) zebrafish larvae were dechorionated and infected at 2 dpf via the yolk sac with wild-type strain of ST, and the number of surviving larvae counted daily during the next 5 days. ST was injected in the yolk sac of 2 dpf *Tg(mpeg1:GAL4)* and *Tg(UAS:DN-asc-GFP)* (C, D, E, F, I) and *Tg(mpx:GAL4)* and *Tg(UAS:DN-asc-GFP)* (G, H, J) larvae and in the otic vesicle of 2 dpf Tg(mpeg1:GAL4:mfap4:tomato) and *Tg(UAS:DN-asc-GFP)* (K, L), *Tg(mpx:GAL4:lyz:DsRED2)* and *Tg(UAS:DN-asc-GFP)* (M, N) larvae. The transcript levels of the indicated genes (C, D, E, F, G, H) were analyzed at 6, 12 or 24 hpi by RT-qPCR in larval head and tail. Macrophage (K, L) and neutrophil (M, N) number and recruitment were analyzed at 1, 3, and 6 hpi by fluorescence microscopy. Caspase-1 activity was determined at 24 hpi using a fluorogenic substrate (I, J). Each dot represents one individual, and the means ± SEM for each group is also shown. P values were calculated using one-way analysis of variance (ANOVA) and Tukey multiple range test. *P ≤ 0.05, **P ≤ 0.01, ***P ≤ 0.001, and ****P ≤ 0.0001. auf, arbitrary units of fluorescence.

We next examined immune cell recruitment and their total number by injecting ST in the otic vesicle. On the one hand, while inhibition of macrophage inflammasome did not affect the recruitment of these cells (Figures 2K and S2A), the total number of macrophages increased in basal condition and upon ST infection at 1 hpi (Figures 2L and S2B). At later timepoints, however, macrophage inflammasome inhibition led to decreased macrophage number, probably reflecting their killing by ST (Figures 2L and S2B). On the other hand, neutrophil recruitment in the larvae with neutrophil inflammasome inhibition turned out to be potentiated as more neutrophils were recruited to the otic vesicle (Figures 2M and S2C). What is more, the total number of neutrophils increased upon inhibition of their inflammasome in basal condition and upon infection at all timepoints analyzed. However, neutrophil counts decreased in wild type larvae upon infection at 3 and 6 hpi (Figures 2N and S2D). These results suggest that ST promoted neutrophil killing through pyroptosis, while other cell death mechanisms should operate in macrophages.

### Overactivation of neutrophil inflammasome leads to myeloid cell imbalance and attenuated inflammation

We next examined the impact of activation of either the neutrophil or the macrophage inflammasome. To do so, we generated the *Tg(lyz:Dendra-Asc)* which overexpressed Dendra-Asc driven by the neutrophil specific promoter *lyz* and additionally a green heart marker. We found that overactivation of neutrophil inflammasome resulted in increased larval resistance to bacterial infection (Figure 3A) and decreased inflammation at 24 hpi than their wild type siblings (Figures 3B-3C). These results were further confirmed using the Nfkb reporter line *Tg(nfkb:GFP)* where expression of Dendra-Asc in neutrophils robustly diminished ST-induced Nfkb reporter activity at 6 and 24 hpi (Figures 3D-3E and S3A-S3B). As expected, caspase-1 activity levels increased in *Tg(lyz:Dendra-Asc)* under basal and infection conditions, confirming the overactivation of the inflammasome (Figure 3F). Furthermore, larvae overexpressing Dendra-Asc in neutrophils exhibited reduced recruitment to ST at 1, 3, and 6 hpi compared to their control siblings (Figures 3G and S3C), likely due to their lower neutrophil numbers in baseline and upon infection at all analyzed timepoints (Figures 3H and S3D). As activation of the inflammasome may lead to pyroptosis, we used TUNEL assay combined with immunostaining of neutrophils with an anti-Mpx antibody. The results confirmed that neutrophil inflammasome overactivation led to higher number of TUNEL positive cells, lower number of neutrophils and higher percentage of TUNEL^+^ neutrophils than their wild-type siblings (Figures 3I-3L and S3E). We then analyzed the number of macrophages in *Tg(lyz:Dendra-Asc)* larvae and we observed higher number of macrophages than control larvae, which explains their more robust recruitment to the bacterium (Figures 3M-3N and S3F-S3G). Strikingly, macrophages of *Tg(lyz:Dendra-Asc)* larvae were resistant to ST killing (Figures 3M-3N and S3F-S3G). Collectively, these results uncover a previously unrecognized crosstalk between neutrophils and macrophages mediated by the inflammasome and further suggest an anti-inflammatory role of macrophages in zebrafish larvae.

**Figure 3.**
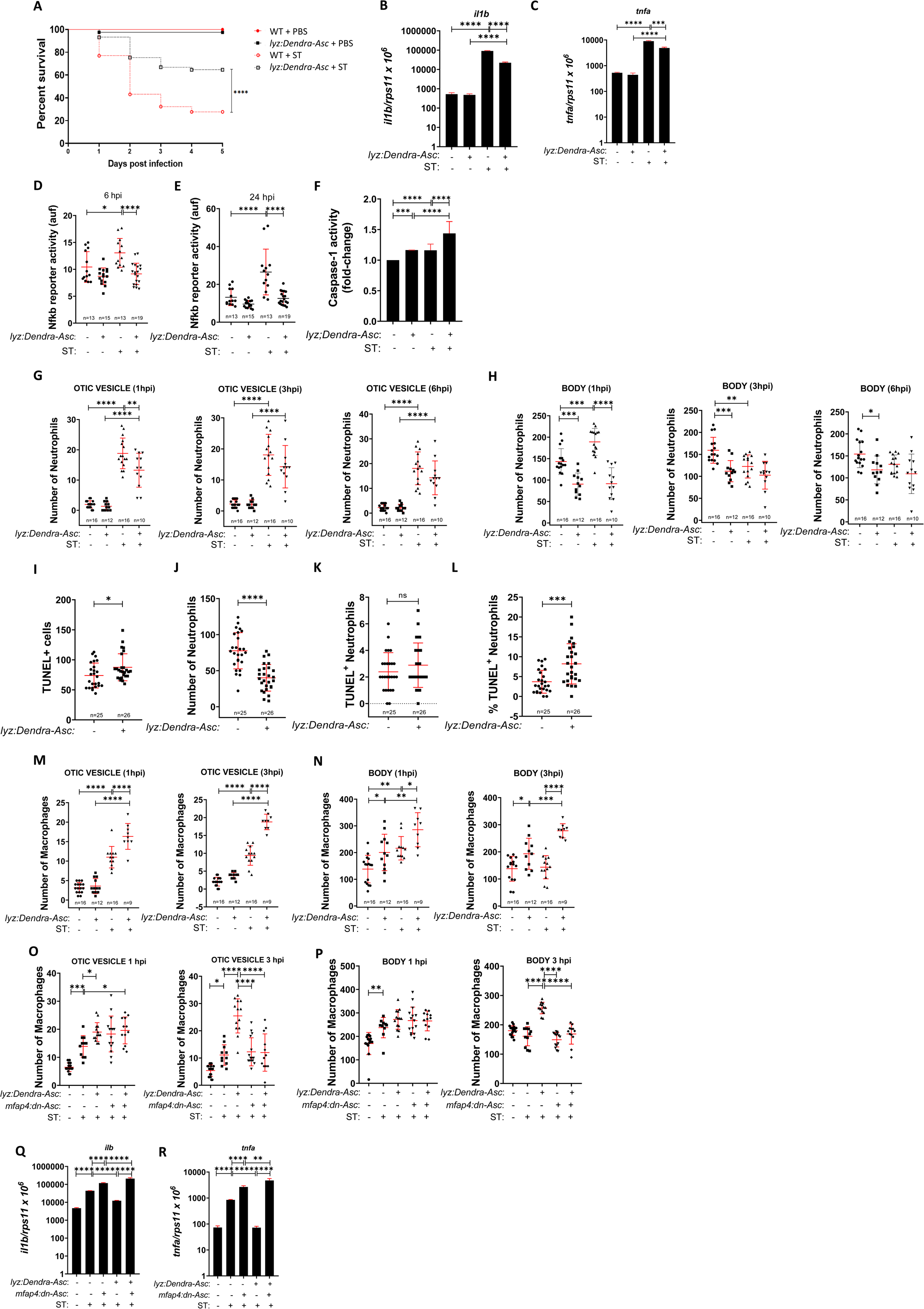
Overactivation of neutrophil inflammasome leads to myeloid cell imbalance and attenuated inflammation. *Tg(lyz:Dendra-Asc)* (A, B, C, F) and *Tg(lyz:Dendra-Asc)* and *Tg(nfkb:eGFP)* (D, E), *Tg(lyz:Dendra-Asc)* and *Tg(lyz:DsRED2)* (G, H), *Tg(lyz:Dendra-Asc)* and *Tg(mfap4:tomato)* (M, N), *Tg(lyz:Dendra-Asc)* and *Tg(mpx:eGFP)* (I, J, K, L) were used. Embryos were infected in yolk sac (A, B, C, D, E, F Q, R) or in the otic vesicle with wild-type strain of ST (G, H, M, N, O, P). Zebrafish one-cell embryos were injected with *mfap4:DN-Asc* plasmid (O, P, Q, R). The number of surviving larvae counted daily during the next 5 days were analyzed (A). The transcript levels of the indicated genes (B, C, Q, R) were analyzed at 24 hpi by RT-qPCR in whole larvae. NfkB activation (D, E) was analyzed at 6 and 24 hpi using a fluorescence microscope. Neutrophil (G, H) and macrophage (M, N, O, P) number and recruitment were analyzed at 1, 3 and 6 hpi by fluorescence microscopy. TUNEL assay and immunohistochemistry staining (I, J, K, L) were performed. Caspase-1 activity was determined at 24 hpi using a fluorogenic substrate (F). Each dot represents one individual, and the means ± SEM for each group is also shown. P values were calculated using Studet’s t-test or one-way analysis of variance (ANOVA) and Tukey multiple range test. *P ≤ 0.05, **P ≤ 0.01, ***P ≤ 0.001, and ****P ≤ 0.0001. auf, arbitrary units of fluorescence.

To further understand the effects of overactivation of the inflammasome in neutrophils and the crosstalk between neutrophils and macrophages, we combined overactivation of inflammasome in neutrophils using *Tg(lyz:Dendra-Asc)* together with inhibition of the macrophage inflammasome expressing DN-Asc in macrophages. The results showed that macrophage inflammasome inhibition abrogated the resistance of macrophages of *Tg(lyz:Dendra-Asc)* to ST killing and the attenuated inflammatory response (Figures 3O-3R and S3H-S3I). This suggests that crosstalk between neutrophil and macrophage inflammasomes is crucial for regulating the inflammatory response, with the macrophage inflammasome playing a previously unrecognized role in the resolution of inflammation.

Similar results were obtained with the COVID-19-associated CSS. Thus, the transcript levels of *il1b* and *tnfa* were significantly lower in *Tg(lyz:Dendra-Asc)* larvae with than in their wild-type siblings upon S1WT injection (Figures S4A-S4D). *Tg(lyz:Dendra-Asc)* larvae showed higher caspase-1 activity than wild-type animals in both basal and upon S1WT injection condition (Figures S4E-S4F). Finally, the immune cell behavior in larvae with overactivated neutrophil inflammasome at 6 hpi was like the one seen with bacterial infection: lower recruitment of neutrophils to the site of the injection and lower number of neutrophils than in control larvae (Figure S4G-S4I and S5A-S5B). However, although *Tg(lyz:Dendra-Asc)* larvae showed normal recruitment of macrophages, they had much more macrophages in basal conditions and upon S1WT injection (Figure S4J-S4L and S5C-S5D).

### Overactivation of the macrophage inflammasome leads to myeloid cell imbalance and enhanced inflammation

The phenotype observed following cell-specific activation of the inflammasome in neutrophils prompted us to investigate the impact of macrophage inflammasome activation. To this end, we microinjected the plasmid *mpeg1:Dendra-Asc* into one-cell stage embryos to achieve transient expression of Dendra-Asc in macrophages. Attempts to generate a stable line were unsuccessful, likely because overactivation of Dendra-Asc in macrophages may be lethal to the fish. Unexpectedly, overactivation of macrophage inflammasome resulted in increased larval susceptibility to ST infection (Figure 4A). Moreover, *mpeg1:Dendra-Asc* larvae showed significantly higher inflammation in both basal and infected conditions than their wild-type siblings, as assayed with the transcript levels of *il1b* and *tnfa* by RT-qPCR at 24 hpi (Figures 4B-4C). These results were further confirmed with the Nfkb reporter line *Tg(nfkb:eGFP)*, where expression of Dendra-Asc in macrophages dramatically potentiated the levels of Nfkb reporter activity in both uninfected and infected larvae at both 6 and 24 hpi (Figures 4D-4E and S6A-S6B). As expected, overexpression of Dendra-Asc in macrophages led to increased caspase-1 activity (Figure 4F).

**Figure 4.**
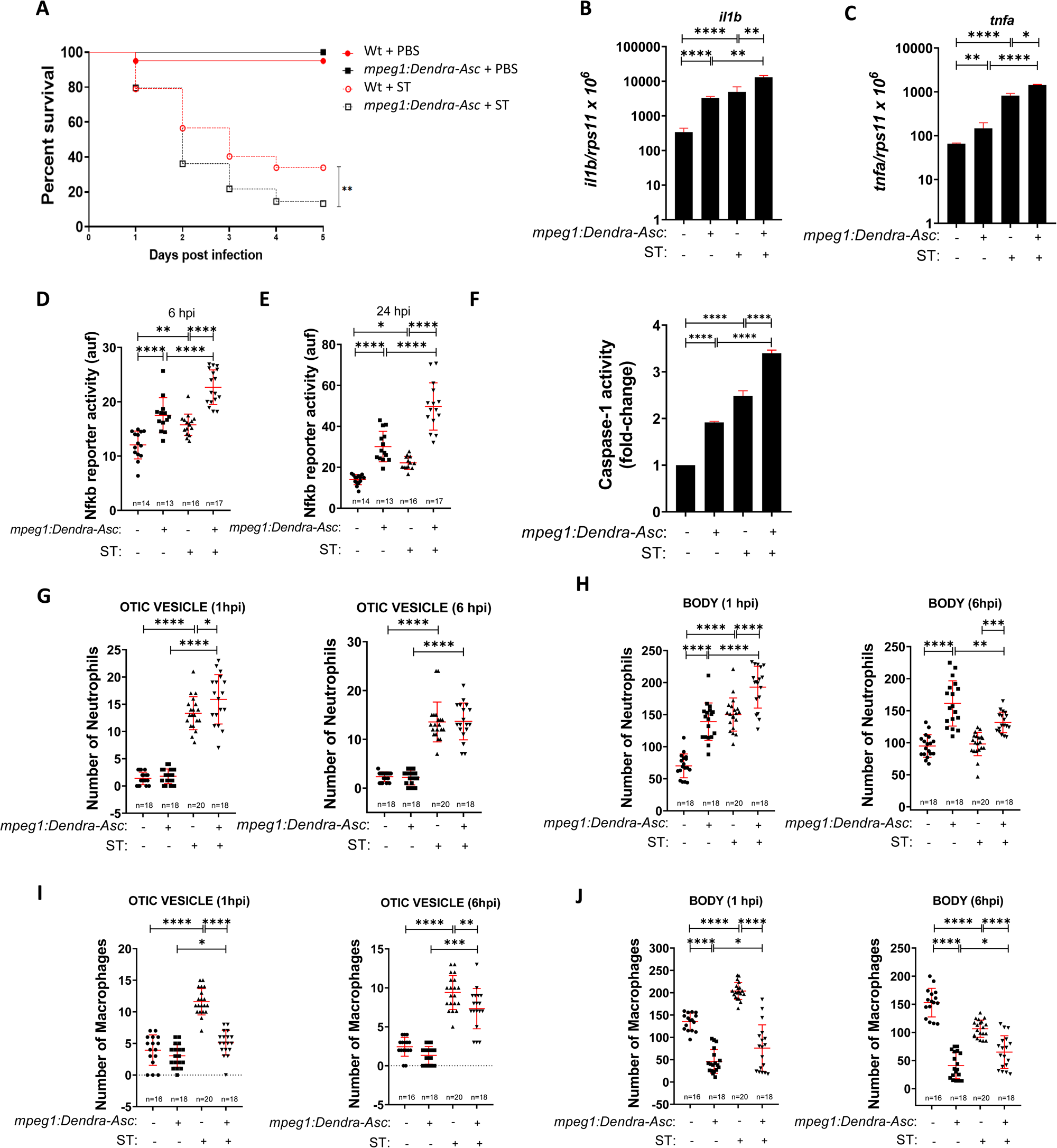
Overactivation of the macrophage inflammasome leads to myeloid cell imbalance and enhanced inflammation in ST infection model. Zebrafish one-cell embryos were injected with *mpeg1:Dendra-Asc* plasmid (A-J). ST was then injected in the yolk sac of 2dpf wild-type larvae (A, B, C, F), *Tg(nfkb:eGFP)* (D, E), or in the otic vesicle of *Tg(mpx:eGFP)* (G, H) and *Tg(mpeg1:mCherry)* (I, J) larvae. The number of surviving larvae were analyzed daily during the next 5 days (A). The transcript levels of the indicated genes (B, C) were analyzed at 24 hpi by RT-qPCR. NfkB activation was analyzed at 6 and 24 hpi (D, E). Caspase-1 activity was determined at 24 hpi using a fluorogenic substrate (F). Neutrophil (G, H) and macrophage (I, J) number and recruitment were analyzed at 1 and 6 hpi by fluorescence microscopy. Each dot represents one individual, and the means ± SEM for each group is also shown. P values were calculated using one-way analysis of variance (ANOVA) and Tukey multiple range test. *P ≤ 0.05, **P ≤ 0.01, ***P ≤ 0.001, and ****P ≤ 0.0001. auf, arbitrary units of fluorescence.

We next analyzed neutrophil and macrophage behaviors in larvae with overactivation of the macrophage inflammasome in response to infection. Stronger recruitment of neutrophils to the bacteria in larvae expressing Dendra-Asc in macrophages than in control larvae was found only at 1 hpi (Figures 4G and S6C-S6D). This may be explained by the neutrophilia found in *mpeg1:Dendra-Asc* uninfected and infected larvae and their killing by ST at 3 (Figures 4H and S6E-S6F). In sharp contrast, the recruitment of macrophages in *mpeg1:Dendra-Asc* larvae decreased in uninfected and infected conditions, probably because of the monocytopenia observed in *mpeg1:Dendra-Asc* injected larvae (Figures 4I-4J and S6G-S6J). Therefore, cell-specific overactivation of the inflammasome led to pyroptotic cell death of both neutrophils and macrophages.

### Extracellular ASC specks drive emergency myelopoiesis and hyperinflammation via inflammasome activation

These unexpected results, showing a crosstalk between neutrophil and macrophage inflammasomes, together with the reported ability of extracellular ASC specks to propagate inflammation (*27, 28*), led us to study the impact of injecting human recombinant ASC protein in circulation of 2 dpf larvae. The results showed increased number of neutrophils and macrophages at 1, 3, 6 and 24 hpi (Figures 5A-5B). The injection itself, as well as the injection of BSA protein as a control, did not cause any increase in the number of neutrophils or macrophages at any time point analyzed (Figure 5A-5B). Moreover, hASC injected larvae showed increased levels of inflammation, namely higher transcript levels of *il1b* and *tnfa* at 3 and 24 hpi than the larvae injected with BSA (Figure 5C). Curiously, inhibition of the canonical inflammasome with the caspase-1 inhibitor VX-765 resulted in a dual effect: rapid attenuation and slow potentiation of hASC-driven induction of *il1b* and *tnfa* (Figure 5C).

**Figure 5.**
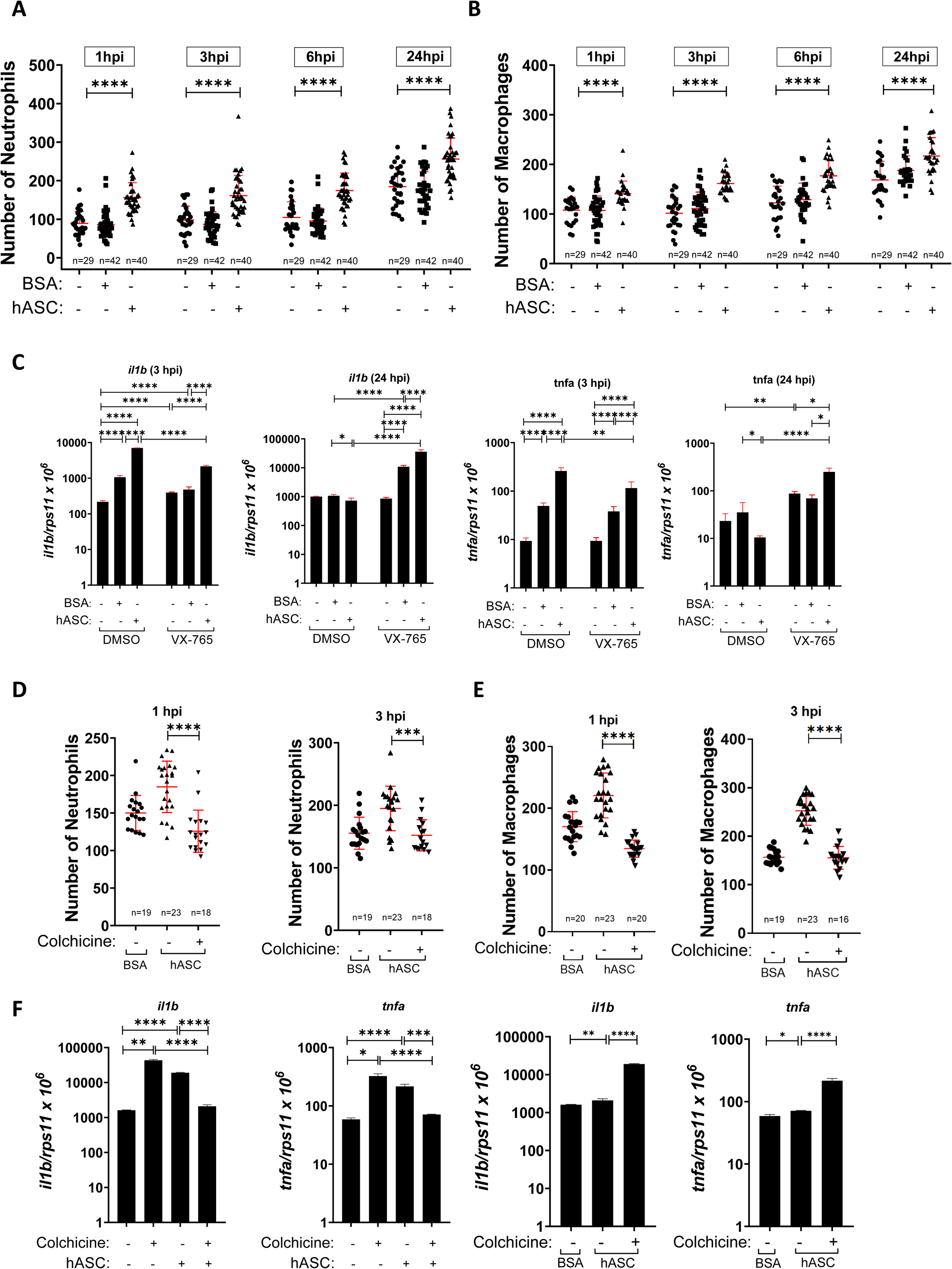
Extracellular ASC specks drive emergency myelopoiesis and hyperinflammation via inflammasome activation. hASC and BSA (as control) were injected in the duct of Cuvier of wild-type (C, F), *Tg(mpx:eGFP)* (A, D) and *Tg(mfap4:tomato)* (B, E) 2 dpf-larvae. In some experiments larvae were pre-treated for 24 h by immersion with DMSO/water, as controls, or with a specific inhibitor of caspase-1 (VX-765, 100 µM) (C) or 1 mg/ml colchicine (D, E, F). The transcript levels of the indicated genes (C, F) were analyzed at 24 hpi by RT-qPCR. Neutrophil (A, D) and macrophage (B, E) number and their recruitment were analyzed at 1, 3, 6 and 24 hpi by fluorescence microscopy. Each dot represents one individual, and the means ± SEM for each group is also shown. P values were calculated using one-way analysis of variance (ANOVA) and Tukey multiple range test. *P ≤ 0.05, **P ≤ 0.01, ***P ≤ 0.001, and ****P ≤ 0.0001. auf, arbitrary units of fluorescence.

Finally, to confirm the role of phagocytosis of hASC specks by zebrafish myeloid cells, wild-type larvae were treated with colchicine and then injected with hASC. The results showed that colchicine was able to block hASC-induced neutrophilia, monocytosis and hyperinflammation (Figures 5D-5F and S7A-S7B). Collectively, these results suggest that hASC specks are taken up by neutrophils and macrophages, activating their inflammasomes to drive emergency myelopoiesis and inflammation. Furthermore, they further point out a dual role of the inflammasome in regulating inflammation.

### Neutrophil to macrophage Asc speck transfer occurs in vivo

We next examined using the unique advantage of the zebrafish model for *in vivo* imaging whether Asc specks can be transferred from neutrophil to macrophage by using photoconvertible green to red fluorescent Dendra (*33*). We first examined Dendra-Asc in *Tg(lyz:Dendra-Asc)* with spinning disk confocal microscopy and found that it was expressed in neutrophils but with two different patterns: either uniformly distributed within the cell or forming Asc specks (Figure 6A). This suggests that overexpression of Dendra-Asc may induce Asc oligomerization without further stimulus, confirming our previous functional and caspase-1 activity results. We next photoconverted Dendra-Asc in the HBV of 2 dpf larvae and we imaged this area for green and red fluorescence 4.5 hours later, confirming photoconversion in several neutrophils (Figure 6B). This photoconversion approach was used in the COVID-19-associated CSS to image neutrophils present at the CHT at 4.5 hpi of S1WT-injected larvae. Green neutrophils with and without red Asc-specks were clearly visible (Figure 6C and Movies 1 and 3), suggesting that activated neutrophils wandered from HBV to CHT. Of note, while the total number of green neutrophils in the CHT was similar between control and S1WT-injected larvae, the S1WT-injected group showed a significantly higher number of neutrophils containing red fluorescent Asc-specks compared to their control siblings (Figure 6D).

**Figure 6.**
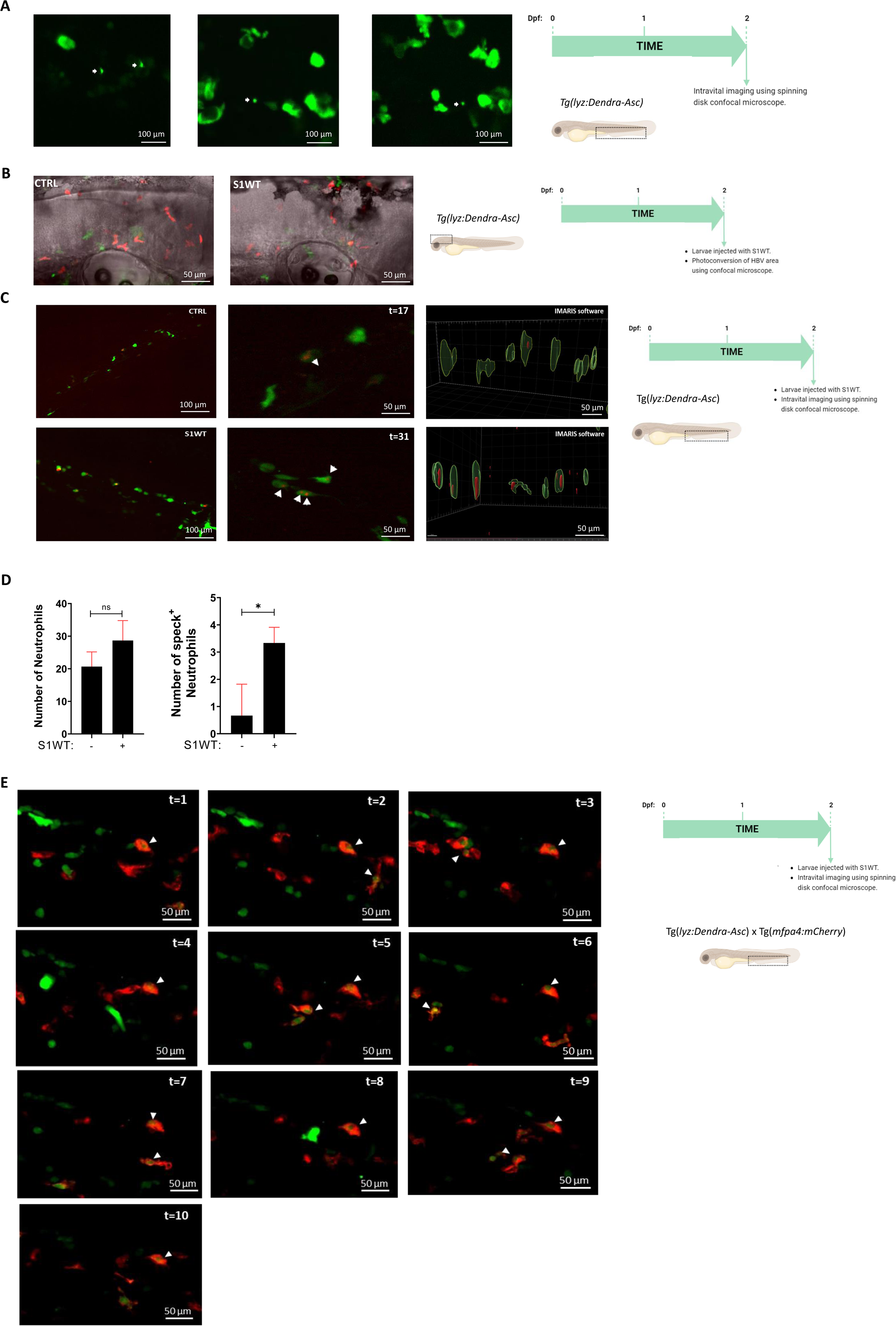
Neutrophil to macrophage Asc speck transfer occurs *in vivo*. Recombinant S1WT (+) or vehicle (−) were injected in the hindbrain ventricle (HBV) of 2 dpf Tg(lyz:Dendra-Asc) (A-D) or *Tg(lyz:Dendra-Asc)*; *Tg(mfap4:mCherry*) (E). HBV photoconversion was performed after injection using LSM880 airyscan (B-D) and high resolution intravital imaging of CHT was performed for 8 h with spinning disk confocal microscopy to visualize *in vivo* Asc-speck formation, release and spreading. 3D IMARIS software was used for image analysis (C). Neutrophil were visualized in green and Asc-speck in red (C, E), while Asc-speck were visualized in green and macrophage in red (E). The total number of neutrophils and the number of neutrophils with speck in S1-injected larvae are shown (D). P values were calculated using Student’s *t*-test. *P ≤ 0.05. The regions of interest used for quantification in all experiments are indicated in the schematic representation of the larvae. Bars: 100 µm (CHT) and 50 µm (HBV).

This result prompted us to examine whether Asc-specks may be transferred from neutrophils to macrophages using *Tg(lyz:Dendra-Asc)*; *Tg(mfap4:mCherry-F)* line. Larvae of 2 dpf were injected with S1WT in the CHT to increase the formation of Dendra-Asc specks in neutrophils and Dendra-Asc speck transfer from neutrophils to macrophages was successfully observed at 4.5 hpi (Figure 6E). Thus, numerous red fluorescent macrophages were observed with green Dendra-Asc specks that originated in Dendra-Asc neutrophils (Figure 6E and Movie 3). Collectively, these results that Asc specks can be transferred from activated neutrophils to bystander neutrophils and from activated neutrophils to bystander macrophages.

### Gsdme- and Ninj1-driven cell death mediate the crosstalk between neutrophils and macrophages

To study whether cell death is involved in the crosstalk between the inflammasome of neutrophils and macrophages, we inactivated the 2 paralogs of zebrafish Gasdermin E (Gsdmea and Gsdmeb), since GSDMD is only present in mammals (*34*) and Gsdme operates downstream the inflammasome in zebrafish (*35, 36*). Gsmdea/b deficiency (edition efficiency of about 60%) fully rescued the neutropenia of Tg(*lyz:Dendra-Asc*) larvae (Figures 7A and S7A) confirming that overactivation of the neutrophil inflammasome resulted in Gsdmea/b-mediated pyroptosis of these cells. Strikingly, the opposite effect was seen with macrophages, where Gsdmea/b deficiency was able to reverse the monocytosis of Tg(*lyz:Dendra-Asc*) larvae to the levels of the negative siblings (Figures 7B and S7B). In addition, Gsdmea/b deficiency was able to rescue the low transcript levels of *il1b* and *tnfa* in larvae with overactivated neutrophil inflammasome (Figure 7C, 7D). Similarly, depletion of Gdsmea/b was able to rescue both monocytopenia and neutrophilia of larvae transiently expressing *mpeg1:Dendra-Asc* (Figures 7E, 7F and S7C, S7D). In sharp contrast, the increased transcript levels of *il1b* and *tnfa* of larvae with overactivated macrophage inflammasome was unaffected by Gsdmea/b deficiency (Figure 7G, 7H). Together, these results highlight the involvement of Gsdmea/b-mediated pyroptosis in the crosstalk between neutrophils and macrophages and reveal that macrophage Gsdme acts as a negative regulator of inflammation.

**Figure 7.**
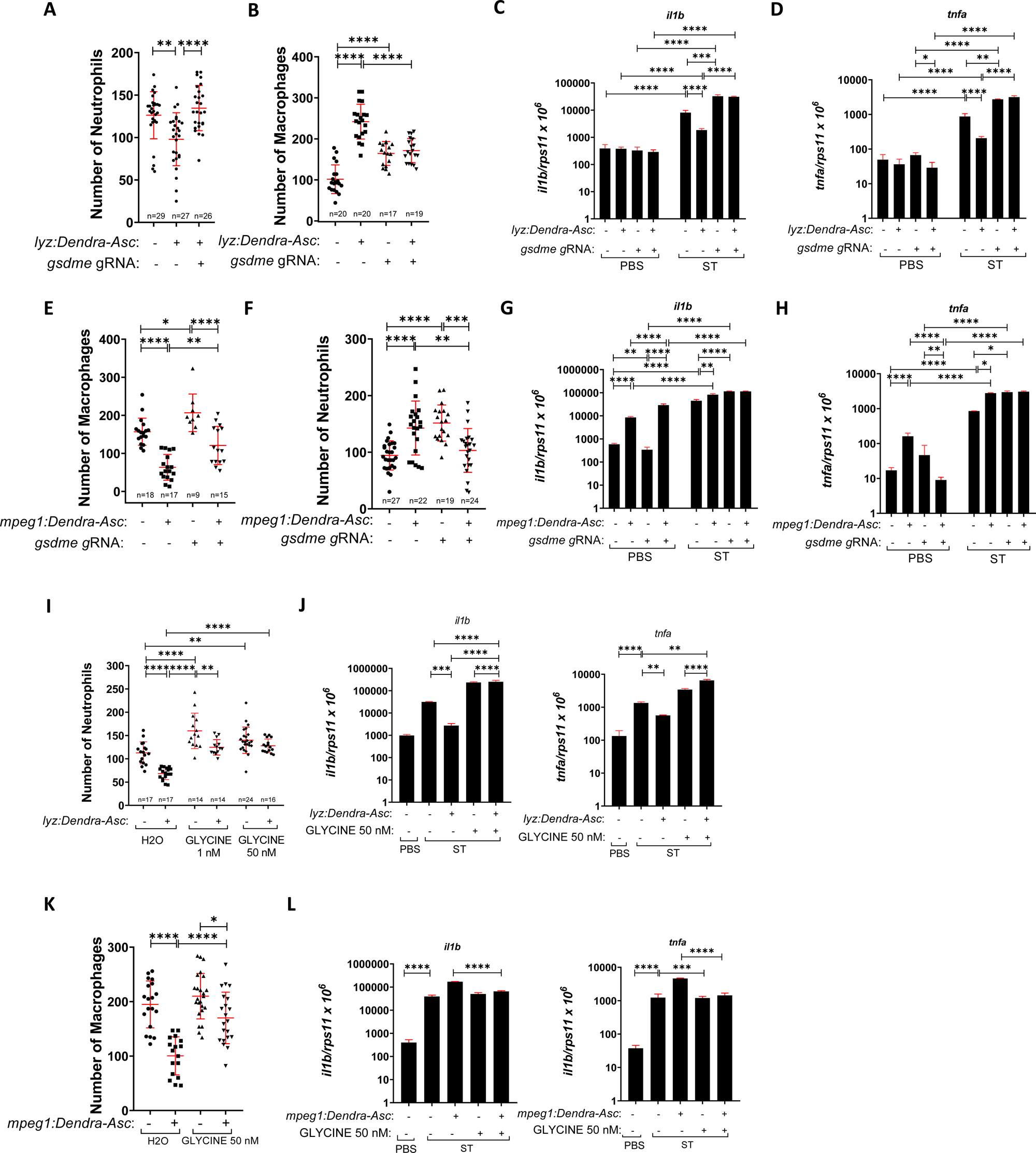
Gsdme- and Ninj1-driven cell death mediate the crosstalk between neutrophils and macrophages. Zebrafish one-cell embryos were injected with *gsdme* gRNA/Cas9 complexes (A-H) or/and with *mpeg1:Dendra-Asc* plasmid (E, F, G, H, K, L). ST was then injected in the yolk sac of 2 dpf *Tg(lyz:Dendra-Asc)* (C, D, J), and *mpeg1:Dendra-Asc* injected (G, H, L) larvae. Zebrafish larvae were also treated by immersion with DMSO or with the specific PMR inhibitor glycine (I, J, K, L). The transcript levels of the indicated genes (C, D, G, H, J, L) were analyzed at 24 hpi by RT-qPCR. Neutrophil (A, F, I) and macrophage (B, E, K) number was analyzed at 2 dpf by fluorescence microscopy. Each dot represents one individual, and the means ± SEM for each group is also shown. P values were calculated using one-way analysis of variance (ANOVA) and Tukey multiple range test. *P ≤ 0.05, **P ≤ 0.01, ***P ≤ 0.001, and ****P ≤ 0.0001.

To confirm the above results, we incubated 1 dpf *Tg(lyz:Dendra-Asc)* larvae during 48 h with glycine, a plasma membrane rupture (PMR) inhibitor (*37, 38*) and found that glycine was able to fully rescue the pyroptotic cell death of neutrophils in *Tg(lyz:Dendra-Asc)* larvae and increased the total number of neutrophils in their wild-type siblings (Figures 7I and S7E). Importantly, glycine also reversed the decreased inflammation of infected *Tg(lyz:Dendra-Asc)* larvae (Figures 7J). Similarly, glycine rescued the pyroptotic cell death of macrophages (Figures 7K and S7F) and decreased the hyperinflammation induced by ST infection (Figures 7L). The same results were obtained by knocking down Ninjurin-1 (Ninj1, KO efficiency of 80%) (Figure S8A-S8F), confirming that Ninj1-driven PMR is required for the crosstalk between neutrophils and macrophages.

We further investigated the role played by Gsdme in the anti-inflammatory function of macrophages by overexpressing catalytic-dead Caspa-C270A, which should behave as a DN form. It was found that while hASC injection resulted in hyperinflammation in Gsdme-deficient larvae, confirming the results with ST infection, expression of Caspa-C270A in neutrophils fully abrogated the pro-inflammatory activity of hASC in both wild type and Gsdme-deficient larvae (Figures S9A, S9B). In sharp contrast, the expression of DN-Caspa in macrophages led to hyperinflammation independently of Gsdme (Figures S9C, S9D). Strikingly, pharmacological inhibition of caspase-1 activity using VX-765 was ineffective in reducing the hyperinflammation and heightened caspase-1 activity in Gsdme-deficient larvae (Figures S9E, S9F). However, VX-765 treatment cooperatively abrogated with Nlrp3 deficiency (KO efficiency of 75%)(*35*) hASC-induced inflammation and caspase-1 activity (Figures S9G, S9H). Finally, genetic inhibition of Gsdme in Nlrp3-deficient larvae phenocopied the effects of VX-765 and Gsdme deficiency (Figures S9I, S9J). These findings underscore the pivotal role of the macrophage Nlrp3 inflammasome and Gsdme in the feedback inhibition of inflammasome activity and the regulation of inflammation.

### The C-terminal fragment of Gsdme negatively regulates inflammasome activation

We next investigated the anti-inflammatory role of Gsdme by overexpressing different fragments of Gsdmea in larvae, since Gsdmea is expressed at higher levels in macrophages than Gsdmeb in zebrafish larvae (*39*). Overexpression of N-terminal (NT) Gsdmea increased the transcript levels of *il1b* and *tnfa* and caspase-1 activity in infected (Figures 8A-8C) and hASC-injected (Figures 8D-8F) larvae. In contrast, C-terminal (CT) Gsdmea had the opposite effects (Figures 8A-8F) in these two models of inflammation. Since these results indicate that CT-Gsdme mediates feedback inhibition of the inflammasome, in contrast to previous findings in mice and humans showing that NT-GSDMD inhibits CASP1 (*40*), we analyzed the activity of full-length wild-type Gsdmea and an uncleavable mutant (Gsdmea-D256A) in the ST infection model. The results showed that while wild-type Gsdmea had minimal impact on inflammation, as indicated by the transcript levels of *il1b* and *tnfa*, and only weakly increased caspase-1 activity, the uncleavable Gsdmea-D256A mutant robustly enhanced both inflammation and caspase-1 activity (Figures 8G-8I). Collectively, these findings demonstrate that the CT fragment of Gsdme plays a critical role in feedback inhibition of inflammasome activity, thereby facilitating the resolution of inflammation.

**Figure 8.**
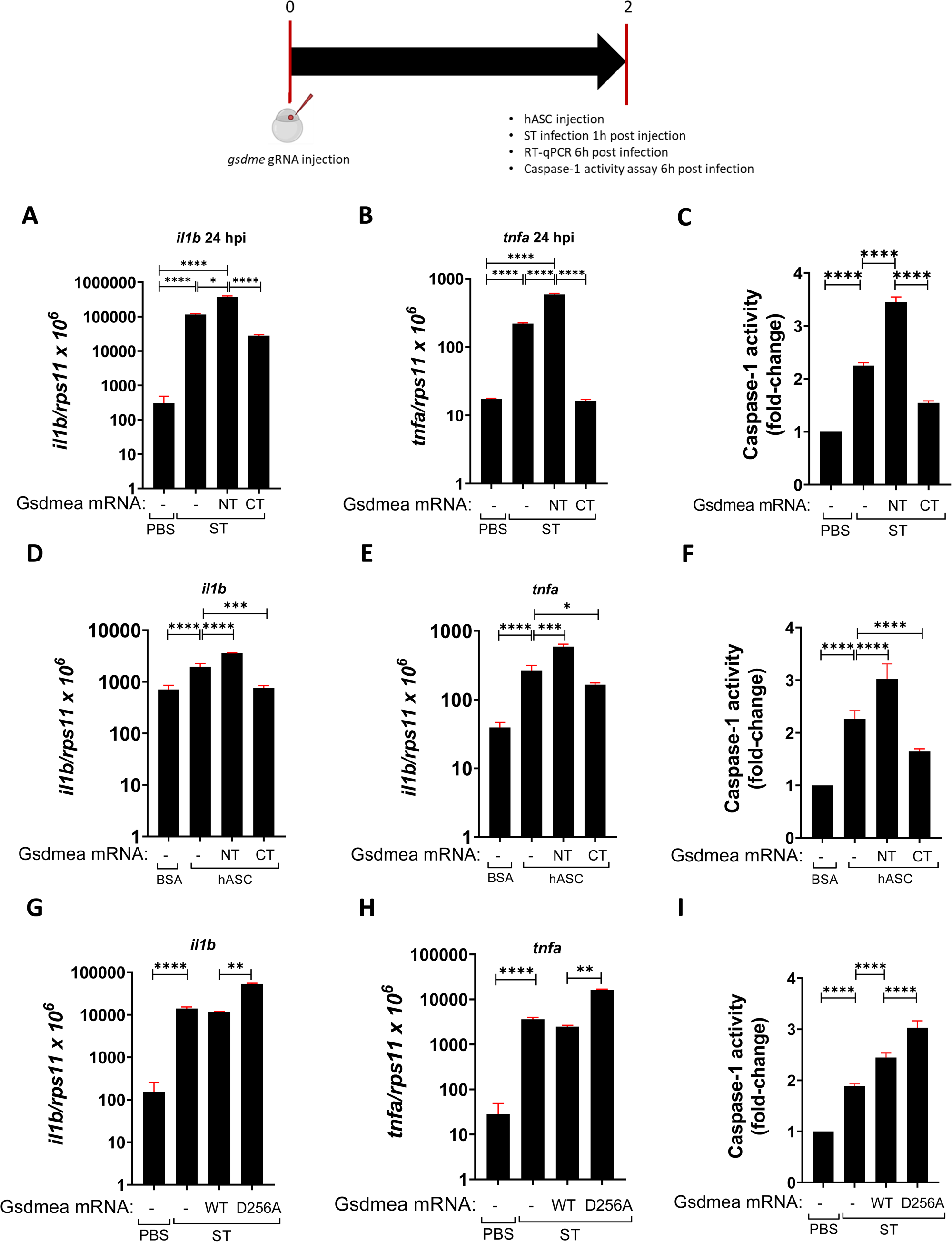
The C-terminal fragment of Gsdme negatively regulates inflammasome activation. Wild-type zebrafish one-cell embryos were injected with mRNA encoding either Gsdmea-NT or Gsdmea-CT mRNA (A-F) or WT or D256A forms of Gsdmea mRNA (G-I). At 2 dpf, larvae were infected with ST via the yolk sac (A-C, G-I) or injected with BSA or hASC via the duct of Cuvier (D-F). The transcript levels of the indicated genes (A, B, D, E, G, H) were analyzed at 24 hpi by RT-qPCR, while caspase-1 activity was also determined at 24 hpi using a fluorogenic substrate (C, F, I). The data are shown as the mean ± SEM. P values were calculated using one-way analysis of variance (ANOVA) and Tukey multiple range test. *P ≤ 0.05, **P ≤ 0.01, ***P ≤ 0.001, and ****P ≤ 0.0001.

## DISCUSSION

While the pro-inflammatory role of inflammasomes in regulating inflammation and immunity is well-established, their potential anti-inflammatory function remains largely unexplored. The name itself is associated with inflammatory processes leading to the release of cytokines and causing pyroptotic cell death (*2, 41*). Inflammasomes are very potent inflammatory mediators that have always been studied as a set in the sense of the whole organism. Animal models available for inflammasome research are not adapted for the study of cell specific inflammasomes (*42–44*). The lack of animal transparency and the scarcity of animal models specifically deficient for different inflammasome components make it difficult to study this area of research. Due to the growing importance of inflammasomes, their study has been extended to non-mammalian vertebrates such as zebrafish (*45, 46*).

Both macrophages and neutrophils express the inflammasome components, which are ubiquitously present in the various cell types of the immune system. While macrophages and dendritic cells were initially credited with inflammasome activation and IL-1β production, neutrophils were traditionally regarded as recipients of pro-inflammatory cytokines(*16*). However, recent studies have demonstrated that NLRP3, NLRC4 and NLR12 inflammasomes can be activated by neutrophils during bacterial infections (*19, 20*). The activation of well-studied macrophages NLRC4 inflammasome causes pyroptosis, while neutrophil inflammasome activation results in NETosis (*47*). This distinction in neutrophils behavior may be explained by their ASC size and by the presence of multiple ASC specks per cell, whereas macrophages possess single individual specks (*4, 23*). What is more, in neutrophils, inflammasome activation leads to the robust production of IL-1β, while in macrophages, it triggers the release of IL-1β along with other cytokines, including IL-1α and IL18 (*24*). Moreover, in macrophages, caspase-1 is more abundant and is primarily responsible for the production of IL-1β, whereas in neutrophils, IL-1β secretion is mostly mediated by serine proteases (*23*). Additionally, recent study has revealed that neutrophils can also secrete IL-1β via an autophagy-dependent mechanism(*25*). Furthermore, in macrophages caspase-1 cleaves GSDMD and produces N-GSDMD p31 fragments, leading to the formation of plasma membrane pores, IL-1β release and pyroptosis. In contrast, in neutrophils although plasma membrane pores are formed by N-GSDMD p31 fragments, they do not induce pyroptosis (*25*). In turn, neutrophil caspase-4 and caspase-11 are activated via noncanonical inflammasome signaling, resulting in neutrophil GSDMD-induced cell death called NETosis, and the induction of extrusion of host-protective antimicrobial NETs (*16, 21, 22*). Likewise, macrophage GBPs play a crucial role in inflammasome activation, IL-1β production and pyroptosis induction. However, in neutrophils, GBPs only mediate prostaglandin biosynthesis and release (*11, 16, 26*). Finally, macrophages undergo pyroptosis, releasing ASC specks into the extracellular space. In contrast, neutrophils release secretory vesicles containing inflammasome components, which serve to amplify the inflammatory response (*16, 27, 28*).

Here, we show for the first time, using unique advantages of zebrafish model, opposed roles of neutrophil and macrophage inflammasomes in *S*. Typhimurium infection (*11*) and COVID-19-associated CRS (*31*) models, with neutrophils being pro-inflammatory and macrophages being anti-inflammatory. To the best of our knowledge this is the first study showing that the macrophage inflammasome plays an anti-inflammatory role, suggesting that cell-specific inflammasome inhibition may be a more effective therapeutic approach to treat intracellular bacterial infections, COVID-19-associated CRS and even chronic inflammatory diseases. The zebrafish animal models developed here with cell specific Asc loss- and gain-of-functions facilitate the study of cell specific inflammasomes, not only neutrophils and macrophages but also other cell types where the inflammasome play a relevant role.

One important observation from our study is that DN Asc was able to block inflammasome activation in both neutrophils and macrophages leading to hypersusceptibility to *S*. Typhimurium, confirming previous results obtained by knocking-down Asc in all cells of the zebrafish larvae (*11*). Furthermore, these studies also revealed that *S*. Typhimurium killed zebrafish neutrophils by pyroptosis, whereas macrophages were killed by another programmed cell death mechanism that we have recently identified as Ripk1-dependent necroptosis(*48*). Although the contribution of macrophages and neutrophils inflammasome in intracellular bacterial clearance and the relevance of pyroptosis is complex, the crosstalk between these myeloid cells has already been appreciated. For example, *in vivo* mouse models, NLRC4 inflammasome activation induces pyroptotic cell death in macrophages, resulting in the release of intracellular bacteria, including *S*. Typhimurium, *Legionella pneumophila* and *Burkholderia thailandensis*, which are then exposed to uptake and killing by reactive oxygen species in neutrophils (*49*). Furthermore, neutrophils are also the main contributors to IL-1β release in mice (*19*) and eicosanoids in zebrafish (*11*) during *S*. Typhimurium infection, since they are highly resistant to pyroptosis in both species.

Our study has also revealed a previously unappreciated crosstalk between neutrophil and macrophage inflammasomes. Thus, overactivation of neutrophil inflammasome by expressing Dendra-Asc promoted their Gsdme-mediated pyroptotic cell death, although apoptosis and necroptosis were also involved, and an unexpected increase of the number of macrophages in basal conditions. This myeloid cell alteration was further altered in both inflammation models, leading to high bacterial resistance and reduced inflammation. More importantly, overactivation of the neutrophil inflammasome rendered macrophages resistant to *S*. Typhimurium killing and this resistance is mediated by the macrophage inflammasome. This is particularly surprising since inhibition of macrophage inflammasome was unable to protect these cells from bacterial killing in wild-type animals. Conversely, overactivation of macrophage inflammasome resulted in Gsdme-mediated pyroptosis of macrophages and increased number of neutrophils, leading to larval hypersusceptibility to the infection and exacerbated inflammation. Notably, this crosstalk between neutrophils and macrophages is strictly dependent on Gsdme and cell death, since Gsdme deficiency, and genetic and pharmacological (glycine administration) inhibition of Ninj1, which block PMR (*37, 38*), reversed myeloid cell number alterations. Both cell types are known to undergo pyroptosis, however this process has been barely related to inflammation resolution (*50–54*). Most of the studies have already demonstrated the GSDMD, functional homolog of zebrafish Gsdme, proinflammatory properties and an important role in host response to bacterial infection (*55, 56*). Using cell specific expression systems, we have demonstrated for the first time Gsdme-dependent anti-inflammatory properties of macrophages. What is interesting, macrophages show higher expression of Gsdmea comparing with neutrophils, which may be the reason for the anti-inflammatory differences between those two innate immune cells.

Since our results suggest a crosstalk between neutrophil and macrophage inflammasome and Asc specks have been found in the serum of patients with chronic inflammatory diseases and can be taken up by macrophages and activate their inflammasome (*27, 28*), we hypothesized that released Asc specks contribute to the spreading of inflammation. Therefore, we generated Tg(*lyz:Dendra-Asc*) line to track Asc specks *in vivo* using photoconvertible Dendra fluorescent green-to-red protein (*57*). Taking advantage of this line, we were able to visualize for the first time Asc speck formation, spreading and transfer *in vivo* from S1WT-activated neutrophils to bystander neutrophils and macrophages, which further support a role for ASC speck release in the spreading of inflammation in chronic inflammatory and infectious diseases and their roles in the observed crosstalk between neutrophils and macrophages. We also observed that neutrophils expressing Dendra-Asc showed two different phenotypes: uniform distribution of Asc within the cell and cells with already formed Asc specks, confirming that overexpression of Asc alone is able to initiate Asc speck formation and inflammasome activation without further stimuli (*58*). Furthermore, the number of Asc-speck positive neutrophils increased in COVID-19-associated CRS model, promoting Asc specks released and uptake by bystander macrophages. Although additional experiments are needed to confirm that the transfer of Asc specks between neutrophils and macrophages regulates the inflammatory response, the increased inflammation and neutrophilia/monocytosis observed upon injection of recombinant hASC and their inhibition by colchicine strongly support this notion.

The most important discovery of our study is the novel role of macrophage inflammasome in the resolution of inflammation. Although it is well known the crucial role played by macrophages in the resolution of inflammation and their plasticity to polarize to different phenotypes, including anti-inflammatory M2 macrophages, as they scan for danger signals and maintain tissue homeostasis, the role of their inflammasome in this process is unknown (*59–61*). Furthermore, it is clear that neutrophil-macrophage cooperation and direct interactions between them impact on the final outcome of their functions (*62, 63*). Using zebrafish model, we uncovered another piece of complexity to the interactions of these cells (Figure 9): Gsdme/Ninj1-mediated pyroptosis of neutrophil upon ST infection results in the releases of Asc specks into intracellular space, which can be uptaken by bystander macrophages promoting Nlrp3 inflammasome activation, Gsdme processing, feedback inhibition of Caspa by the C-terminal fragment of Gsdme and enhanced anti-inflammatory properties. The anti-inflammatory properties of Gsdme are not unexpected, since mouse GSDMD has been found to play both pro- and anti-inflammatory roles (*64*), and to directly inhibit CASP1 activity (*40*). Although the tetrapeptide within the N-terminal fragment that is required for CASP1 inhibition is relatively well conserved between zebrafish Gsdmea/b and mammalian GSDMD (RFWK in mouse, WFWK in human GSDMD, and WFWQ in zebrafish Gsdmea/b), this conservation does not translate into functional equivalence. Indeed, neither the N-terminal fragment nor the full-length zebrafish Gsdmea were able to inhibit inflammasome activation *in vivo*. In contrast, we found that the C-terminal fragment of zebrafish Gsdmea mediated negative regulation of inflammasome activation in this species. This observation is consistent with the distinct subcellular localization of Gsdme fragments (membrane-associated N-terminal vs. cytosolic C-terminal) and with structural data showing that the CASP1 active site engages not only the GSDMD N- and C-domain linker but also a hydrophobic pocket within the C-terminal domain(*65*). Therefore, our results suggest that the C-terminal fragment of zebrafish Gsdme exerts an additional function as a negative regulator of Caspa activity, beyond its established role in autoinhibition. It remains to be determined whether mammalian GSDMD C-terminal fragments have a similar regulatory function and why Gsdme mediates this anti-inflammatory effect in macrophages but not in neutrophils.

**Figure 9.**
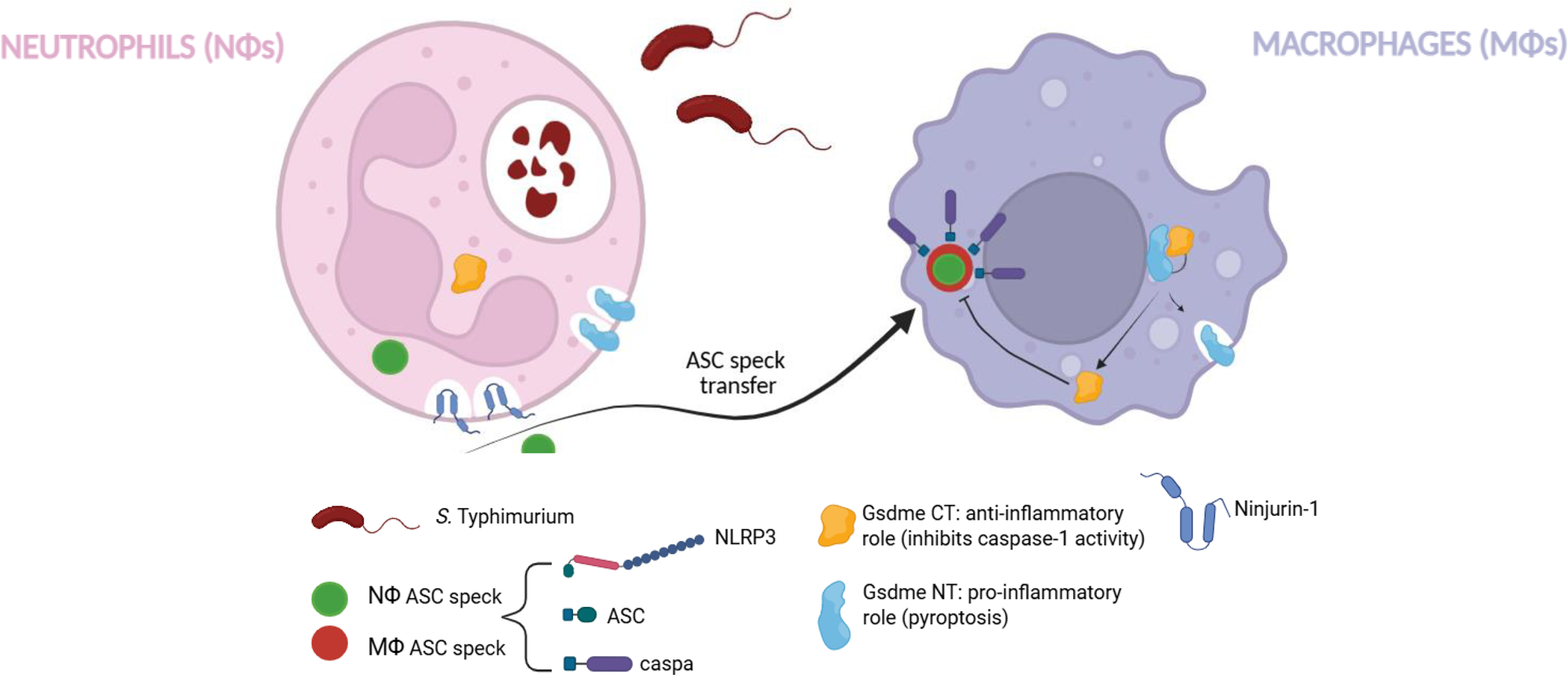
Model of *in vivo A*sc speck transfer from neutrophils to macrophages and its role in inflammasome activation and anti-inflammatory responses. The proposed model illustrates how Gsdme/Ninj1-mediated inflammasome activation in neutrophils during *Salmonella* Typhimurium infection leads to the release of ASC specks into the intracellular space. These specks are taken up by macrophages, promoting Nlrp3 inflammasome activation, Gsdme processing, and feedback inhibition of Caspase-1 activity through the C-terminal fragment of Gsdme. This process is associated with enhanced anti-inflammatory properties in macrophages.

In summary, we present a unique model to study the impact of cell-specific inflammasomes in intracellular bacterial infection and COVID-19-associated CRS. Our findings demonstrate that neutrophil and macrophage inflammasomes play opposing roles in regulating inflammation, highlighting the importance of Gsdme in resolving inflammation, akin to mammalian GSDMD. Furthermore, we provide novel evidence of the transfer of Asc specks from activated neutrophils to bystander neutrophils and macrophages *in vivo*. Understanding the intricacies of this process opens new horizons for developing therapeutic targets to treat inflammasome-dependent infectious and chronic inflammatory diseases.

## MATERIALS AND METHODS

### Animals

Zebrafish (*Danio rerio* H.) were obtained from the Zebrafish International Resource Center and mated, staged, raised and processed as described (*66*). The lines Tg(*lyz:dsRED*)^nz50^ (*67*), Tg(*mpx:eGFP*)^i114^ (*68*), Tg(*mfap4:mCherry-F*)^ump6^ (*69*), Tg(*mfap4:tomato*)^xt12^ (*70*), Tg(*mpeg1:mCherry/tnfa:eGFP-F*)^ump5^ (*71*), Tg(*mpeg1:GAL4*)^gl25^ (*72*), Tg(*mpx:Gal4.VP16*)^i222^ (*73*), Tg(*NFkB-RE:eGFP*)^sh235^ referred to as nfkb:eGFP (*74*), Tg(*UAS:ascΔCARD-GFP*)^ums4^ (*32*), Tg(*UAS:nfsB-mCherry*)^c264^ (*73*), and casper (*mitfaw2/w2; mpv17a9/a9*) (*75*) were previously described. The experiments performed comply with the Guidelines of the European Union Council (Directive 2010/63/EU) and the Spanish RD 53/2013. The experiments and procedures were performed approved by the Bioethical Committees of the University of Murcia (approval number #395/2017). Most experiments were performed with embryos/larvae less than 5 dpf.

### DNA Construct and Generation of Transgenics

The *lyz:Dendra-Asc*, *mpeg1:Dendra-Asc*, *mfap4:dn-Asc* and *uas:dn-caspa* constructs were generated by MultiSite Gateway assemblies using LR Clonase II Plus (Life Technologies) according to standard protocols and using Tol2kit vectors described previously(*76*). The expression construct of Gcsfa(*77*) was previously described. The expression construct of NT-Gsdmea, CT-Gsdmea, full-length wild-type and D256A mutant Gsdmea were synthetized in GeneScript.

Tg(*lyz:Dendra-Asc*) and Tg(*uas:DN-caspa*) were generated by microinjecting 0.5–1 nl into the yolk sac of one-cell-stage embryos a solution containing 50 ng/μl *lyz:Dendra-Asc* construct, respectively, and 150 ng/μl Tol2 RNA in microinjection buffer (× 0.5 Tango buffer and 0.05% phenol red solution) using a microinjector (Narishige).

### CRISPR and RNA injection in zebrafish

crRNA for zebrafish nlrp3, gsdmea, gsdmeb, ninj1 and negative control crRNA (Catalog #1072544), and tracrRNA were purchased from IDT and resuspended in Nuclease-Free Duplex Buffer to 100 µM. 1µl of each was mixed and incubated for 5 min at 95 0C for duplexing. After removing from the heat and cooling to room temperature, 1.43 µl of Nuclease-Free Duplex Buffer was added to the duplex, giving a final concentration of 1000 ng/µl. Finally, the injection mix was prepared by mixing 1 µl of duplex, 2.55 µl of Nuclease-Free Duplex Buffer, 0.25 µl Cas9 Nuclease V3 (IDT, #1081058) and 0.25 µl of phenol red, giving final concentrations of 250 ng/µl of gRNA duplex and 500 ng/µl of Cas9. The prepared mix was microinjected into the yolk of one- to eight-cell-stage embryos using a microinjector (Narishige) (0.5–1 nl per embryo). The same amounts of gRNA were used in all the experimental groups. The efficiency of gRNA was checked by amplifying the target sequence with a specific pair of primers (Table S1) and the TIDE webtool (https://tide.nki.nl/).

In vitro-transcribed RNA was obtained following manufacturer’s instructions (mMESSAGE mMACHINE kit, Ambion). RNA was mixed in microinjection buffer and microinjected into the yolk of one-cell-stage embryos using a microinjector (Narishige; 0.5–1 nl per embryo). The same amount of RNA was used for all the experimental groups.

### Chemical treatments

In some experiments, 1 dpf embryos were manually dechorionated and treated for 1 to 3 dpf at 28°C by bath immersion with Metronidazole (Mtz, 5 mM, Sigma-Aldrich) in egg water supplemented with 1% DMSO. In some experiments, 24 hpf embryos were treated with 0.3% N-Phenylthiourea (PTU, Sigma-Aldrich) to inhibit melanogenesis.

For image acquisition and cell count experiments, 1-day postfertilization (dpf) embryos were manually dechorionated and treated until 48 hpf by bath immersion with Glycine (Sigma-Aldrich, #G7126) at final concentration of 1 or 50 mM diluted in egg water and VX-765 (Belnacasan, MedChemExpress, #HY-13205) at final concentration of 100 μM diluted in egg water, For gene expression experiments, 2 dpf were anesthetized in embryo medium with 0.16 mg/mL tricaine and 10 bacteria/zebrafish were injected into the yolk sac, glycine treatment was refreshed and maintained until 3 dpf (24 hpi) when larvae were collected for further analysis.

Larvae of 2 dpf were treated with 1 mg/ml Colchicine (MedChemExpress, #HY-16569) for 30 min before hASC injection and remained until 1 hpi until *in vivo* imaging was initiated.

### Recombinant protein injection

Recombinant His-tagged Spike S1 wild-type produced in baculovirus-insect cells and with <1.0 EU per μg protein as determined by the LAL method (#40591-V08B1, Sino Biological) at a concentration of 0.25 mg/ml supplemented with phenol red were injected into the hindbrain ventricle (1 nl) of 48 hpf zebrafish larvae (*30, 31*).

Recombinant hASC (#CSB-EP890936HU, Cusabio) was resuspended in sterile water at the concentration of 1 mg/ml. Larvae of 2 dpf were anesthetized in embryo medium with 0.16 mg/mL tricaine and hASC (1 mg/ml), BSA (1 mg/ml) or sterile water was injected into the duct of Cuvier where it is directly drained to blood. The larvae were allowed to recover in egg water at 29°C, and monitored for clinical signs of inflammation.

### Infection assays

For infection experiments, ST 12023 (WT) was used. Overnight cultures in Luria-Bertani (LB) broth were diluted 1/5 in LB with 0.3 M NaCl, incubated at 37 °C until 1.5 optical density at 600 nm was reached, and finally diluted in sterile PBS. Larvae of 2 dpf were anaesthetized in embryo medium with 0.16 mg ml−1 tricaine and 10 bacteria (yolk sac) or 100 (otic vesicle) per larvae were microinjected. Larvae were allowed to recover in egg water at 28–29 °C and monitored for clinical signs of disease or mortality over 5 days. At least three independent experiments were performed with a total number of 200–350 larvae per treatment.

### Analysis of gene expression

Total RNA was extracted from whole larvae or head/tail larvae with TRIzol reagent (Invitrogen) following the manufacturer’s instructions and treated with DNase I, amplification grade (1 U/mg RNA: Invitrogen). SuperScript IV RNase H Reverse Transcriptase (Invitrogen) was used to synthesize first-strand cDNA with random primer from 1µg of total RNA at 50 ^0^C for 50 min. Real-time PCR was performed with an ABIPRISM 7500 instrument (Applied Biosystems) using SYBR Green PCR Core Reagents (Applied Biosystems). Reaction mixtures were incubated for 10 min at 95 ^0^C, followed by 40 cycles of 15 s at 95 ^0^C, 1 min at 60 ^0^C, and finally 15 s at 95 ^0^C, 1 min 60 ^0^C, and 15 s at 95 ^0^C. For each mRNA, gene expression was normalized to the ribosomal protein S11 gene (*rps11*) content in each sample, using the Pfaffl method (*78*). The primers used are shown in Table S2. In all cases, each PCR was performed with triplicate samples and repeated at least with two independent samples.

### Caspase-1 activity assays

The caspase-1 activity was determined with the fluorometric substrate Z-YVAD 7-Amido-4-trifluoromethylcoumarin (Z-YVAD-AFC, caspase-1 substrate VI, Calbiochem), as described previously (*11, 32*). In brief, 25–35 whole larvae or 45 heads/tails were lysed in hypotonic cell lysis buffer (25 mM 4-(2-hydroxyethyl) piperazine-1-ethanesulfonic acid, 5 mM ethylene glycol-bis(2-aminoethylether)-N,N,Ń,Ń-tetraacetic acid, 5 mM dithiothreitol, 1:20 protease inhibitor cocktail (Sigma-Aldrich), pH 7.5) on ice for 10 min. For each reaction, 100 µg protein were incubated for 90 min at room temperature with 50 mM YVAD-AFC and 50 µl of reaction buffer (0.2% 3-[(3-cholamidopropyl)dimethylammonio]-1-propanesulfonate (CHAPS), 0.2 M 4-(2-hydroxyethyl) piperazine-1-ethanesulfonicacid, 20% sucrose, 29 mM dithiothreitol, pH 7.5). After incubation, the fluorescence of the AFC released from the Z-YVAD-AFC substrate was measured with a FLUOstart spectofluorometer (BGM, LabTechnologies) at an excitation wavelength of 405 nm and an emission wavelength of 492 nm. A representative caspase-1 activity graph out of three repeats is shown in figures.

### Imaging of zebrafish larvae

To study immune cell recruitment to the injection site and Nfkb activation, 2 dpf *Tg(mpx:eGFP)*, *Tg(lyz:dsRed)*, *Tg(mfap4:tomato)* or *Tg(nfkb:egfp)* larvae were anaesthetized in embryo medium with 0.16 mg/ml tricaine. Images of the hindbrain, head or the whole-body area were taken 1, 3, 6, 12 and 24 h post-injection (hpi) using a Leica MZ16F fluorescence stereomicroscope. The number of neutrophils or macrophages was determined by counting visually and the fluorescence intensity was obtained and analyzed with ImageJ (FIJI) software (*79*). In all experiments, images were pooled from at least 3 independent experiments performed by two people and using blind samples.

For laser confocal microscopy imaging, larvae were anesthetized in 200 µg/ml tricaine, mounted on 35 mm glass-bottom dishes (WillCo-dish), immobilized in 1% low-melting-point agarose and covered with 2 ml egg water supplemented with 160 µg/ml tricaine. Epi-fluorescent microscopy was performed using a MVX10 Olympus MacroView microscope (MVPLAPO 1X objective and XC50 camera). Confocal microscopy was performed on ANDOR CSU-W1 confocal spinning disk on an inverted NIKON microscope (Ti Eclipse) with ANDOR Neo sCMOS camera (20x air/NA 0.75 objective); laser 488 nm for GFP and 561 nm for mCherry. Image stacks for time-lapse movies were acquired at 28°C. The 4D files generated from time-lapse acquisitions were analyzed using Image J, adjusting brightness and contrast for maximal visibility.

For photoconversion of Dendra protein*, Tg(lyz:Dendra-Asc)* embryos were raised to 2 dpf in the dark, and S1WT-injected larvae in the Hindbrain Ventricle (HBV), as described above, were mounted in 1% low-melting point agarose. A 405-nm Laser Cube 405-50C on a confocal TCS SP5 inverted microscope with HCXPL APO 40x/1.25-0.75 oil objective (Leica) was used to photoconvert the Dendra-labelled cells using 10% laser power (optimized before the experiments). HBVs were imaged before and after the photoconversion in the green (488 nm) and red (561 nm) channels.

### TUNEL assay and immunohistochemistry staining

The number of apoptotic cells in 3 dpf larvae was assessed by TUNEL assay. The larvae were fixed with 4% paraformaldehyde for 2 hours and dehydrated with methanol at −20°C. After gradual rehydration, the embryos were treated with 100% acetone at −20°C for 10 min and rinsed 3 times for 10 min with PBT. Embryos were permeabilized with a solution containing 0.1% Triton X-100 and 0.1% sodium citrate in PBS for 30 minutes at RT and rinsed 2 times for 5 min with PBT. They were then washed 5 times with PBST for 5 minutes each and blocked in 10% calf serum, 1% DMSO, 0.1% Tween-20 for 1 h to initiate whole body immunohistochemistry staining (WIHC).

To label neutrophils, larvae were incubated with anti-Mpx antibody overnight at 4°C (Genetex #GTX128379, 1:150), rinsed 5 times for 5 min with PBST, incubated with a secondary antibody (Invitrogen #A21207, 1:250) for 1 hour and, finally, rinsed 3 times for 5 min with PBST and stored in 70% glycerol in PBS until image acquisition. Images were acquired in a Leica MZ16F fluorescence stereo microscope.

### Neutral Red Staining

Neutral red stains zebrafish macrophages granules and the procedure was performed as originally reported (*80*). Briefly, macrophage staining was performed on live 3dpf larvae and was obtained by the incubation of the embryos in 2.5 g/ml of neutral red solution from Sigma-Aldrich (in embryo medium) at 25–30°C in the dark for 5-8 h. The larvae were anesthetized in 0.16 mg ml-1 tricaine and imaged using a Leica MZ16F fluorescence stereo microscope.

### Statistical analysis

Statistical analysis was performed using Prism 8.0 (GraphPad Software, CA, USA). No calculation was performed to predetermine sample size; experiments were repeated three times to ensure robustness. Embryos were randomly allocated to each experimental condition and analyzed in blind samples. No data were excluded from the analysis. Data are shown as mean ± s.e.m. and were analyzed by analysis of variance and a Tukey multiple range test to determine differences between groups. The differences between the two samples were analyzed by the two-sided Student’s t-test. The data met normal distribution assumption when required and showed similar variances. A log-rank test was used to calculate the statistical differences in the survival of the different experimental groups. A p-value of <0.05 was considered statistically significant.

## ACKNOWLEDGMENTS

We warmly thank I. Fuentes, P. Martínez, M. Rodenas and ME Rubio for their excellent technical assistance. We also thank Profs. S.A. Renshaw, P. Crosier, A.H. Meijer, D. Tobin and L.I. Zon for the zebrafish lines. The authors used ChatGPT (GPT-5, OpenAI) to assist in improving the grammar and language clarity of the manuscript. No content was generated by the tool, and all scientific writing, data interpretation, and conclusions were entirely produced and verified by the authors.

## AUTHOR CONTRIBUTIONS

SDT and VM conceived the study; SDT, AP, JMLG, MENC and GL performed the research; SDT, AP, JMLG, MENC, GL, DGM, MLC and VM analyzed the data; and SDT and VM wrote the manuscript with minor contributions from other authors.

---

**Figure S1 (related to Figure 1).**
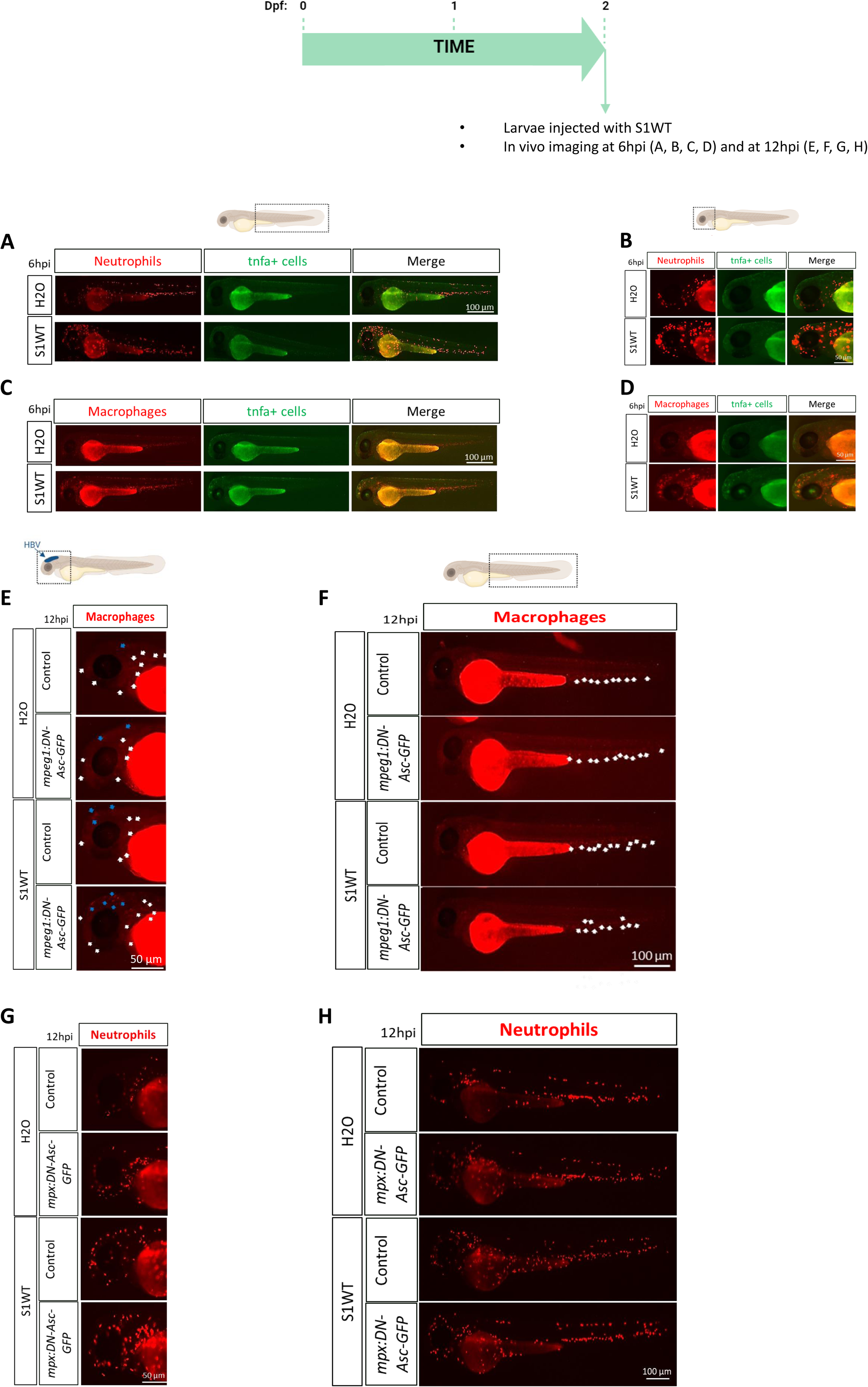
Representative green and red fluorescence images of larvae of the different groups shown in Figure 1. Recombinant S1WT or vehicle (-) were injected in the hindbrain ventricle (HBV) of 2 dpf *Tg(lyz:DsRED2)* and *Tg(tnfa:eGFP)* (A, B), *Tg(mfap4:tomato)* and *Tg(tnfa:eGFP)* (C, D), *Tg(mpeg1:GAL4, mfap4:tomato)* and *Tg(UAS:DN-asc-GFP)* (E, F), *Tg(mpx:GAL4:lyz:DsRED2)* and *Tg(UAS:DN-asc-GFP)* (G, H) larvae. Neutrophils in the body (A, H) and in the HBV and head (B, G), and macrophages in the body (C, F) and in the HBV and head (D, E) visualized at 6 or 12 hpi by fluorescence microscopy. The regions of interest used for quantification in all experiments are indicated in the schematic representation of the larva. Bars: 100 µm (whole larva) and 50 µm (head).

**Figure S2 (related to Figure 2).**
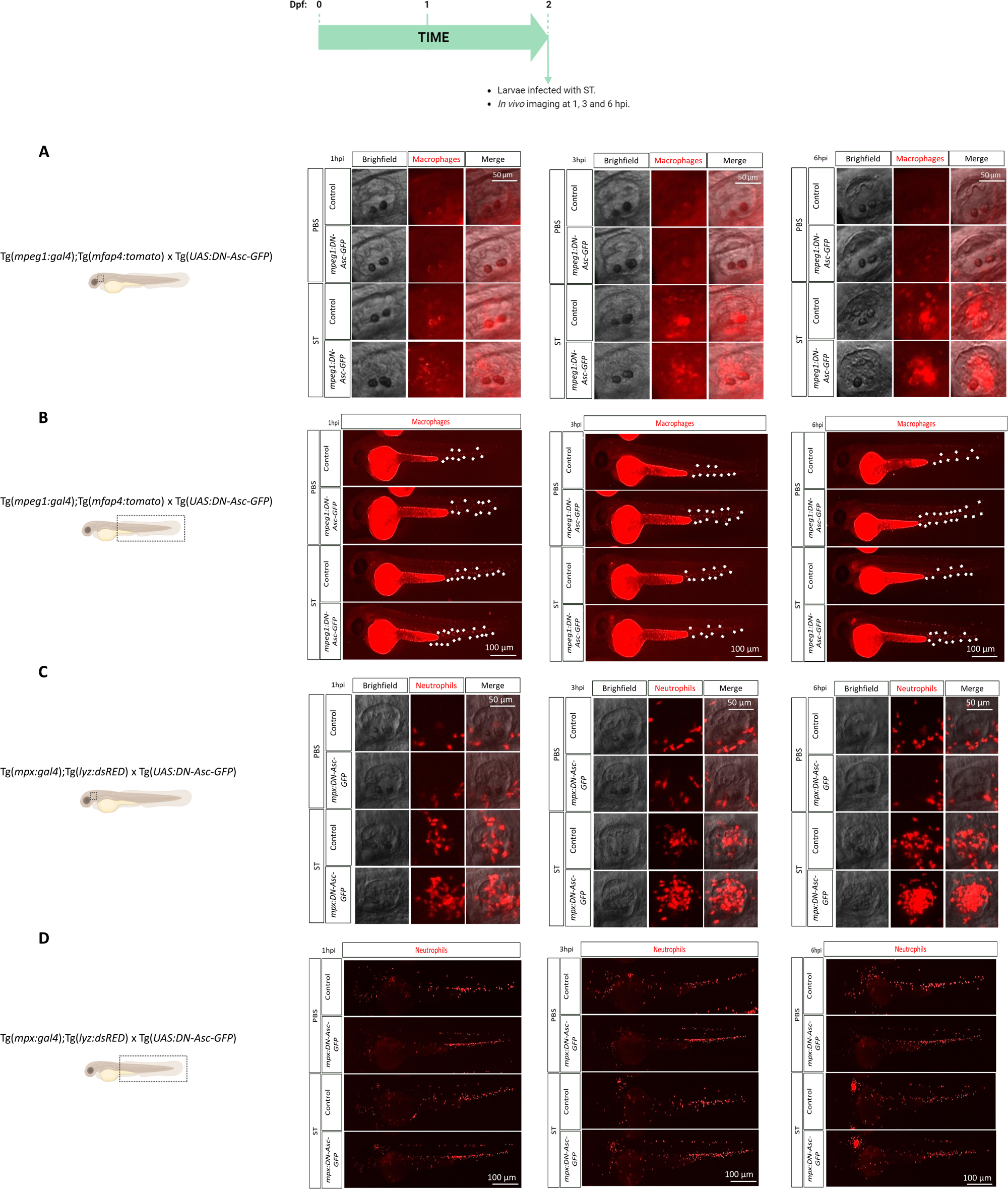
Representative red fluorescent images of larvae of the different groups shown in Figure 2. ST was injected in the otic vesicle of 2 dpf *Tg(mpeg1:GAL4, mfap4:tomato)* and *Tg(UAS:DN-asc-GFP)* (A, B), *Tg(mpx:GAL4:lyz:DsRED2)* and *Tg(UAS:DN-asc-GFP)* (C, D) larvae. Macrophages in the otic vesicle (A) and in the body (B), and neutrophils in the otic vesicle (C) and in the body (D) were visualized at 1, 3 and 6 hpi by fluorescence microscopy. The regions of interest used for quantification in all experiments are indicated in the schematic representation of the larvae. Bars: 100 µm (whole larva) and 50 µm (otic vesicle)..

**Figure S3 (related to Figure 3).**
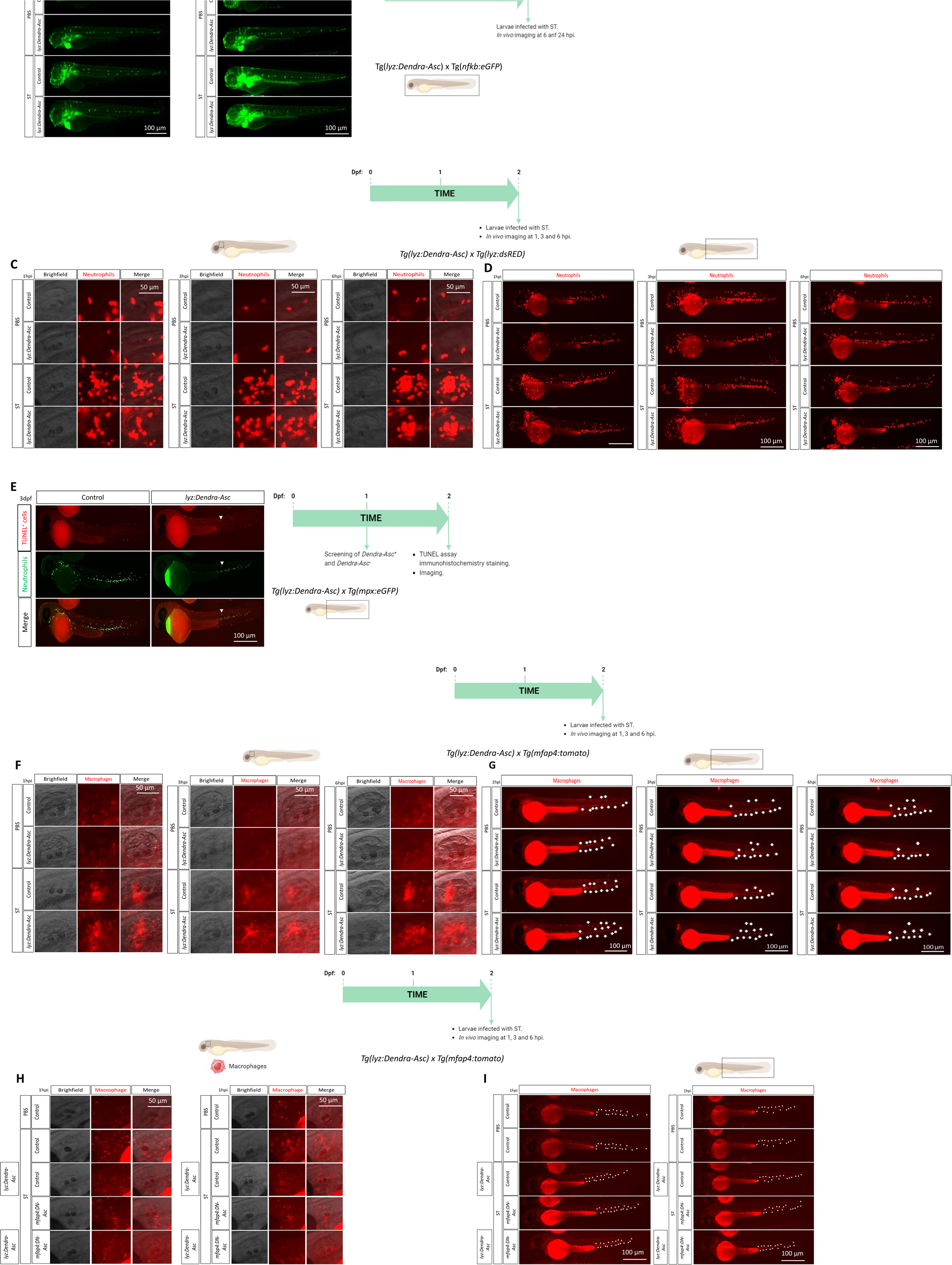
Representative green and red fluorescence images of larvae of the different groups shown in Figure 3. ST was injected in the yolk sac of 2 dpf *Tg(lyz:Dendra-Asc) and Tg(nfkb:eGFP)* (A, B), and in the otic vesicle of 2 dpf *Tg(lyz:Dendra-Asc)* and *Tg(lyz:DsRED2)* (C, D), *Tg(lyz:Dendra-Asc)* and *Tg(mfap4:tomato)* (F, G, H, I), larvae. Neutrophils in the otic vesicle (C) and in the body (D), and macrophages in the otic vesicle (F, H) and in the body (G, I) were visualized at 1, 3 and 6 hpi by fluorescence microscopy. The number of neutrophils and the number of TUNEL positive cells (E) were quantified using a fluorescence microscopy. The regions of interest used for quantitation in all experiments are indicated in the schematic representation of the larvae. Bars: 100 µm (whole larva) and 50 µm (otic vesicle).

**Figure S4 (related to Fig. 3).**
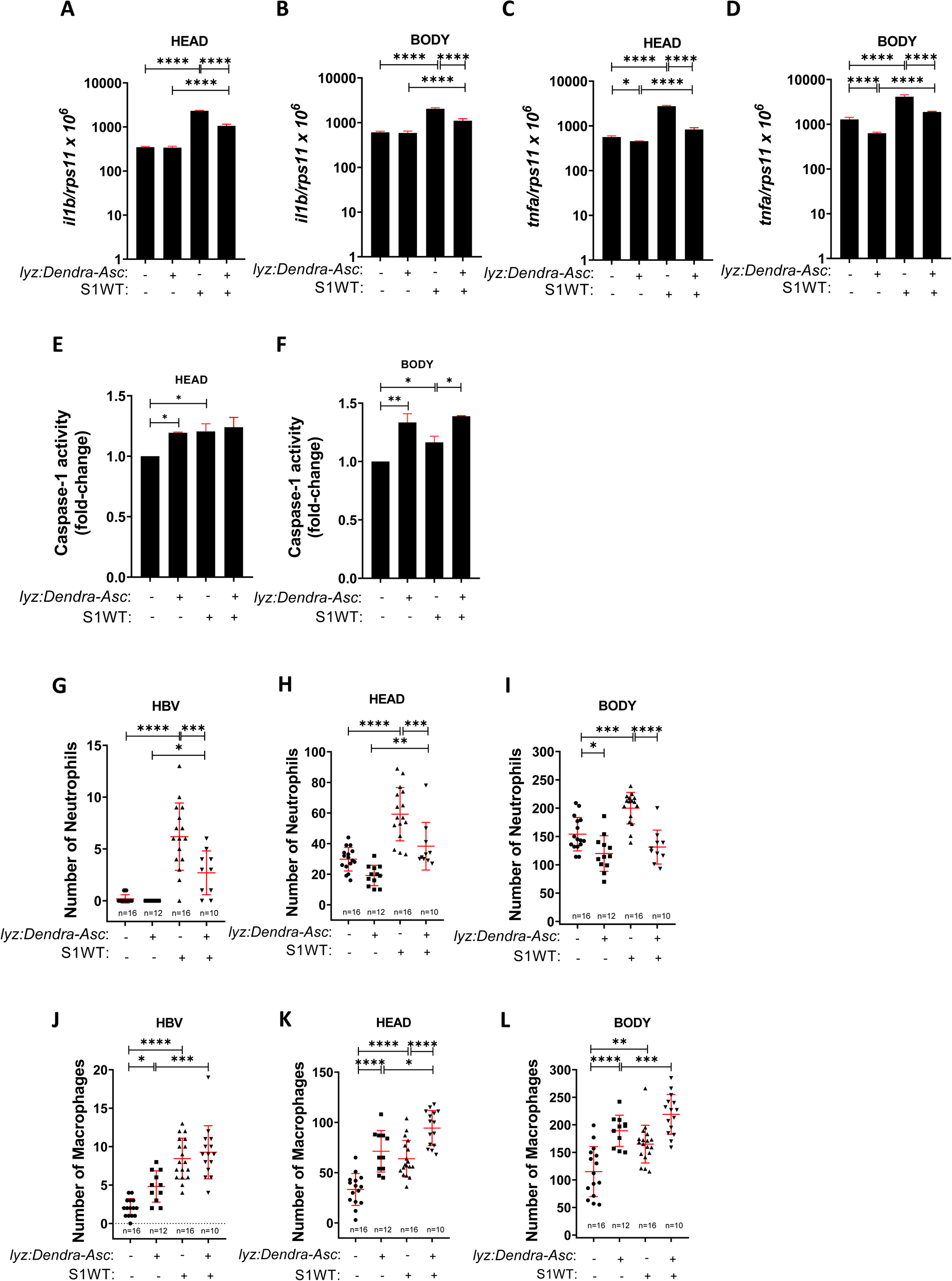
Overactivation of neutrophil inflammasome leads to myeloid cell imbalance and attenuated inflammation. Recombinant S1WT (+) or vehicle (−) were injected in the hindbrain ventricle (HBV) of 2-day postfertilization (dpf) *Tg(lyz:Dendra-Asc)* (A, B, C, D, E, F). *Tg(lyz:Dendra-Asc)* and *Tg(lyz:DsRED2)* (G, H, I), *Tg(lyz:Dendra-Asc*) and *Tg(mfap4:tomato)* (J, K, L) larvae. The transcript levels of the indicated genes (A, B, C, D) were analyzed at 24 hpi by RT-qPCR in larval head and tail. Neutrophil (G, H, I) and macrophage (J, K, L) number and their recruitment were analyzed at 6 hpi by fluorescence microscopy. Caspase-1 activity was determined at 24 hpi using a fluorogenic substrate (E, F). Each dot represents one individual, and the means ± SEM for each group is also shown. P values were calculated using one-way analysis of variance (ANOVA) and Tukey multiple range test. *P ≤ 0.05, **P ≤ 0.01, ***P ≤ 0.001, and ****P ≤ 0.0001. auf, arbitrary units of fluorescence.

**Figure S5 (related to Figure 3).**
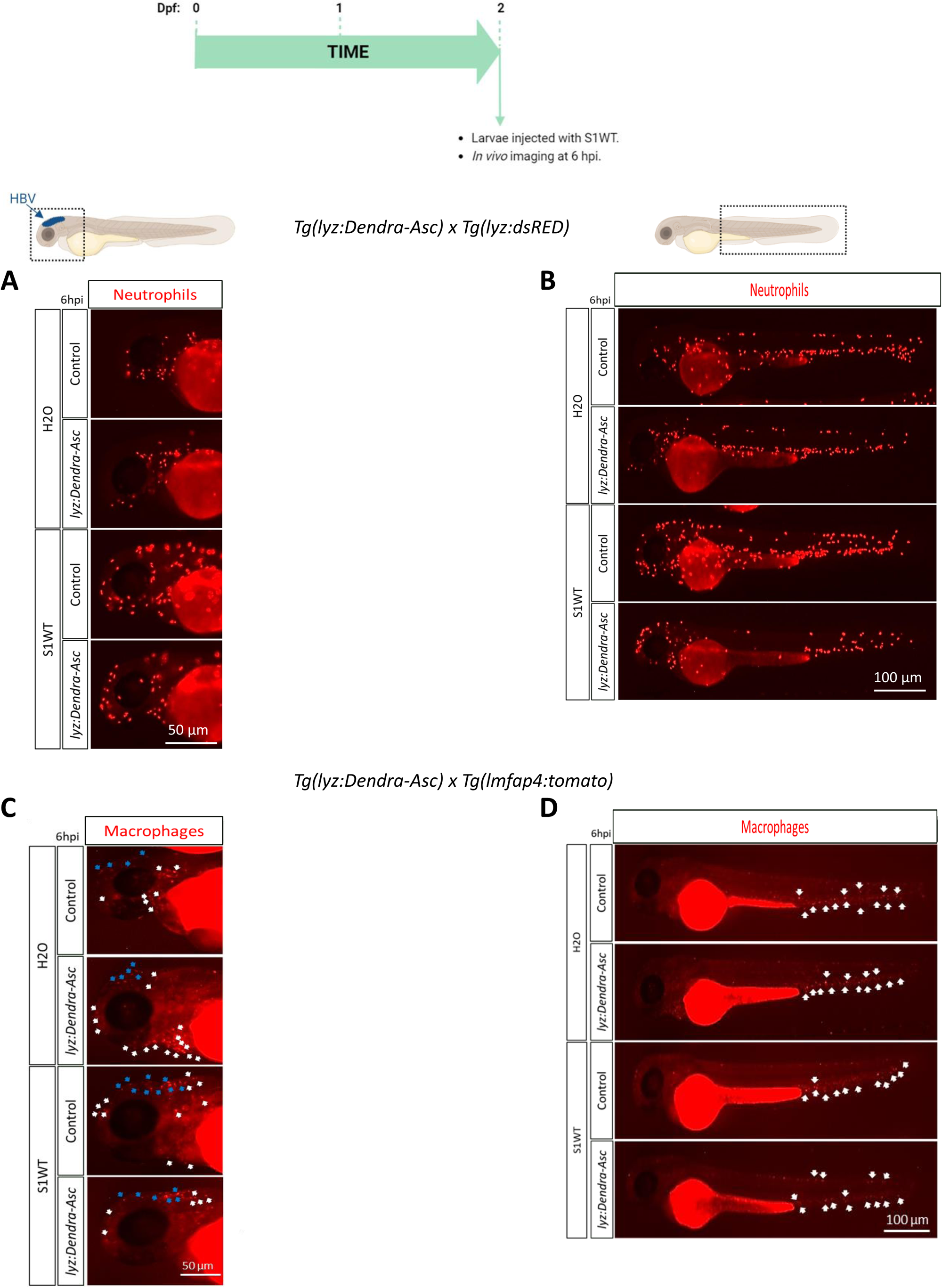
Representative red fluorescence images of larvae of the different groups shown in Figures S4. Recombinant S1WT or vehicle (-) were injected in the hindbrain ventricle (HBV) of 2 dpf *Tg(lyz:Dendra-Asc)* and *Tg(lyz:DsRED2)* (A, B), *Tg(lyz:Dendra-Asc)* and *Tg(mfap4:tomato)* (C, D) larvae. Neutrophils in the head (A) and in the body (B), and macrophages in the head (C) and in the body (D) visualized at 6 hpi by fluorescence microscopy. The regions of interest used for quantification in all experiments are indicated in the schematic representation of the larvae. Bars: 100 µm (whole larva) and 50 µm (head).

**Figure S6 (related to Figure 4).**
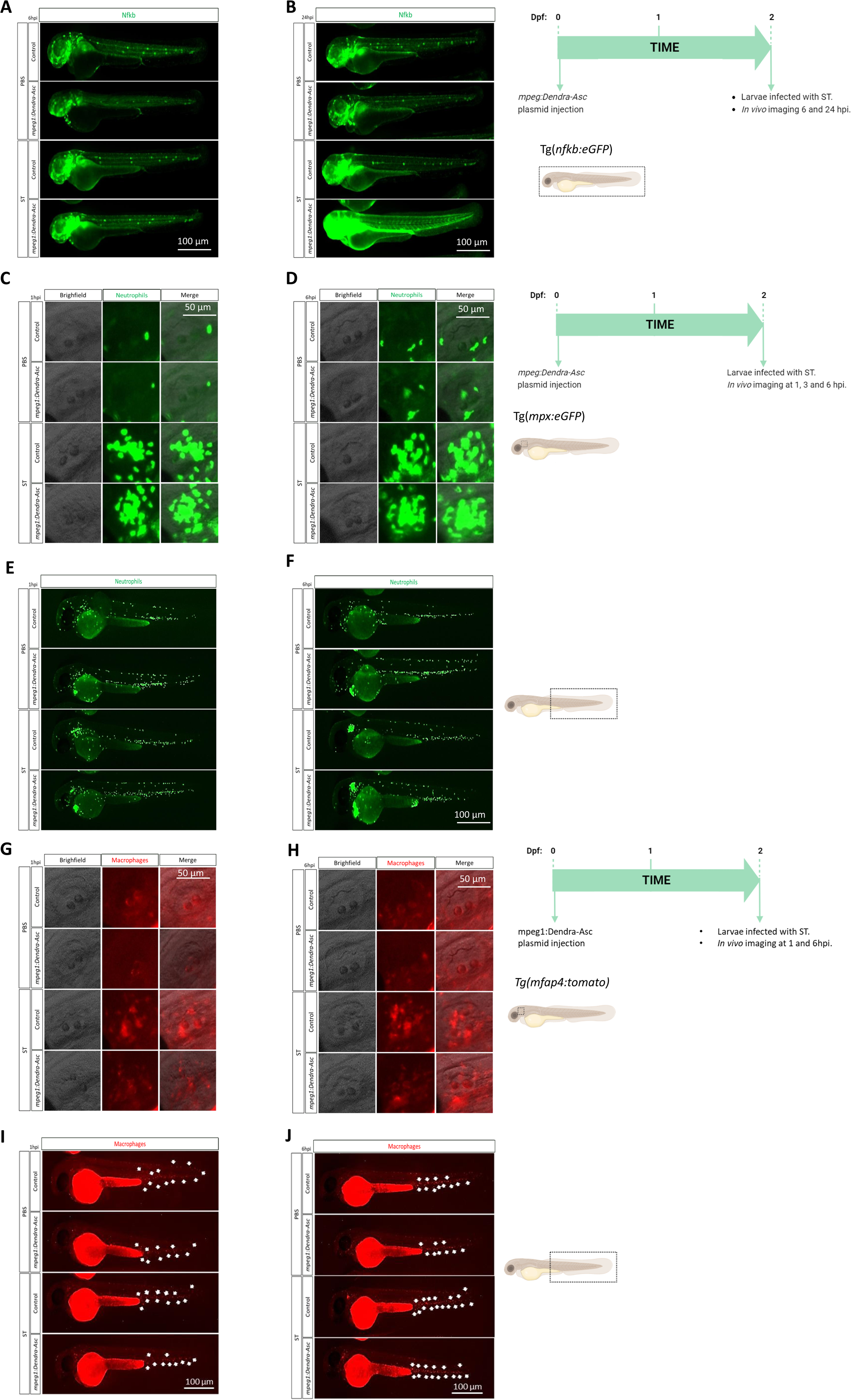
Representative green and red fluorescence images of larvae of the different groups shown in Figures 4. Zebrafish one-cell stage embryos were injected with *mpeg1:Dendra-Asc* plasmid and infected with ST in the yolk sac of 2 dpf *Tg(nfkb:eGFP)* (A, B) and in the otic vesicle of 2 dpf *Tg(mpx:eGFP)* (C, D, E, F), *Tg(mfap4:tomato)* (G, H, I, J) larvae. Nfkb activity (A, B) was visualized at 6 and 24 hpi by fluorescence microscopy. Neutrophils in the otic vesicle (C, D) and in the body (E, F), and macrophages in the otic vesicle (G, H) and in the body (I, J) were visualized at 1 and 6 hpi by fluorescence microscopy. The regions of interest used for quantification in all experiments are indicated in the schematic representation of the larvae. Bars: 100 µm (whole larva) and 50 µm (head).

**Figure S7 (related to Figure 5).**
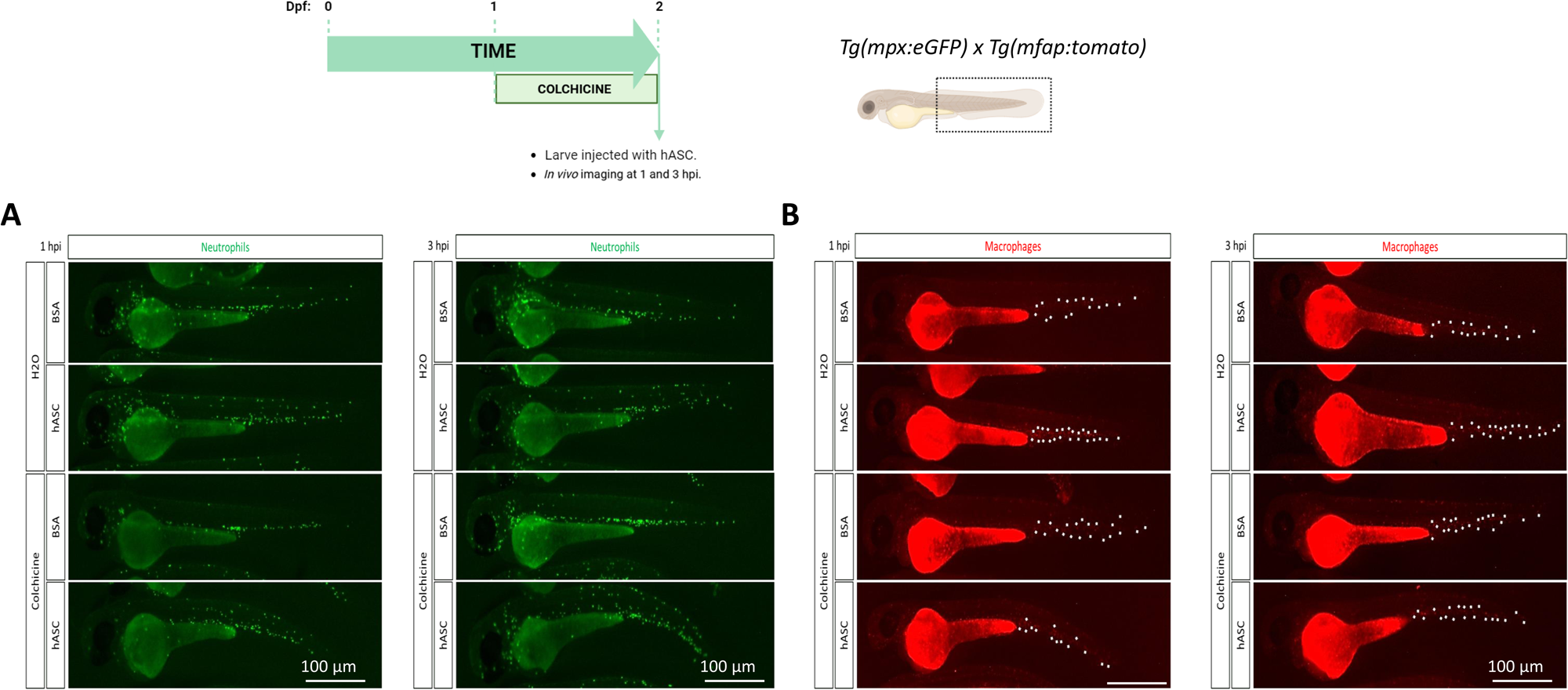
Representative green and red fluorescence images of larvae of the different groups shown in Figure 5. Two-dpf *Tg(mpx:eGFP)* (A) and *Tg(mfap4:tomato)* (B) larvae pre-treated by immersion with 1 mg/ml colchicine for 24h were injected in the duct of Cuvier with either hASC or BSA (control). Neutrophil (A) and macrophage (B) number was analyzed by fluorescence microscopy at 1 and 3 hpi. The regions of interest used for quantification in all experiments are indicated in the schematic representation of the larvae. Bars: 100 µm (whole larva) and 50 µm (head).

**Figure S8 (related to Figure 7).**
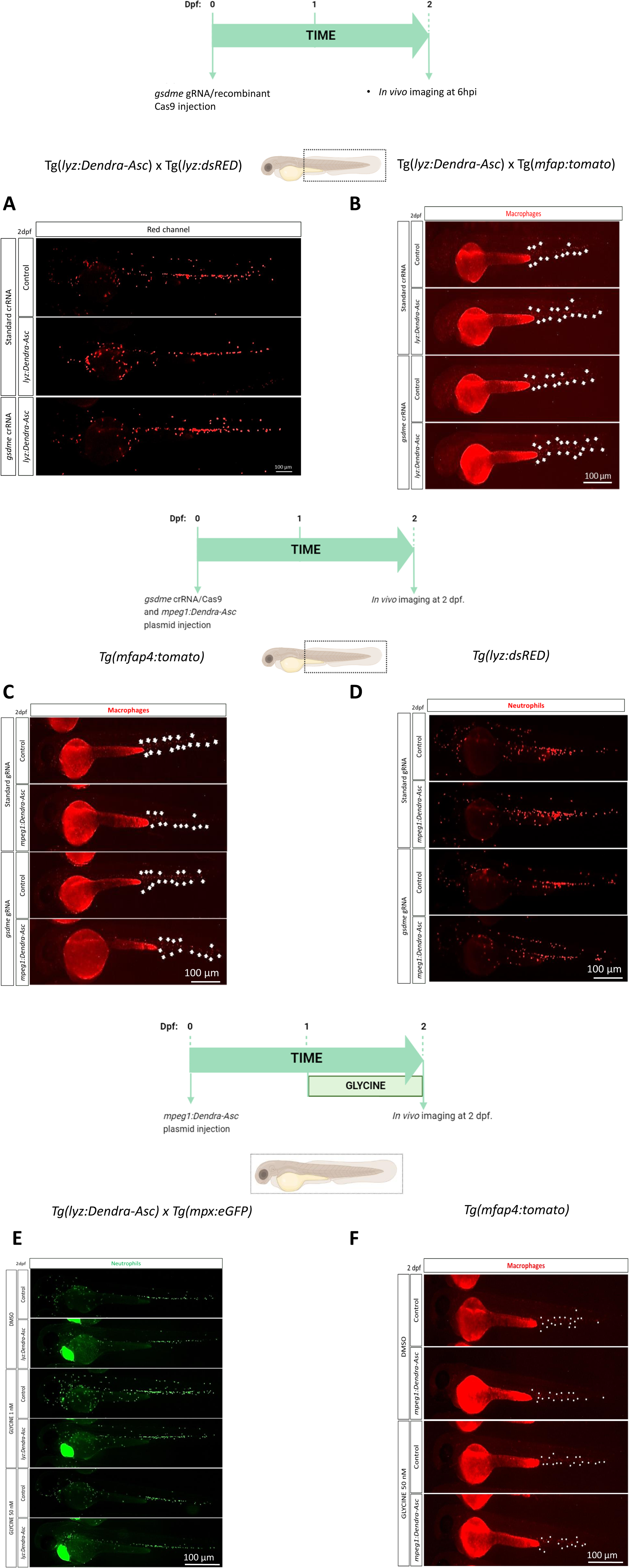
Representative green and red fluorescence images of larvae of the different groups shown in Figure 7. Zebrafish one-cell embryos were injected with gsdme gRNA/Cas9 complexes (A-D) or/and with *mpeg1:Dendra-Asc* plasmid (C, D, F). Zebrafish larvae were also treated by immersion with DMSO or with the specific PMR inhibitor glycine (E, F). Neutrophils (A, D, E) and macrophages (B, C, F) in the body were visualized at 2 dpf by fluorescence microscopy. The regions of interest used for quantification in all experiments are indicated in the schematic representation of the larvae. Bars: 100 µm.

**Figure S9 (related to Fig. 7).**
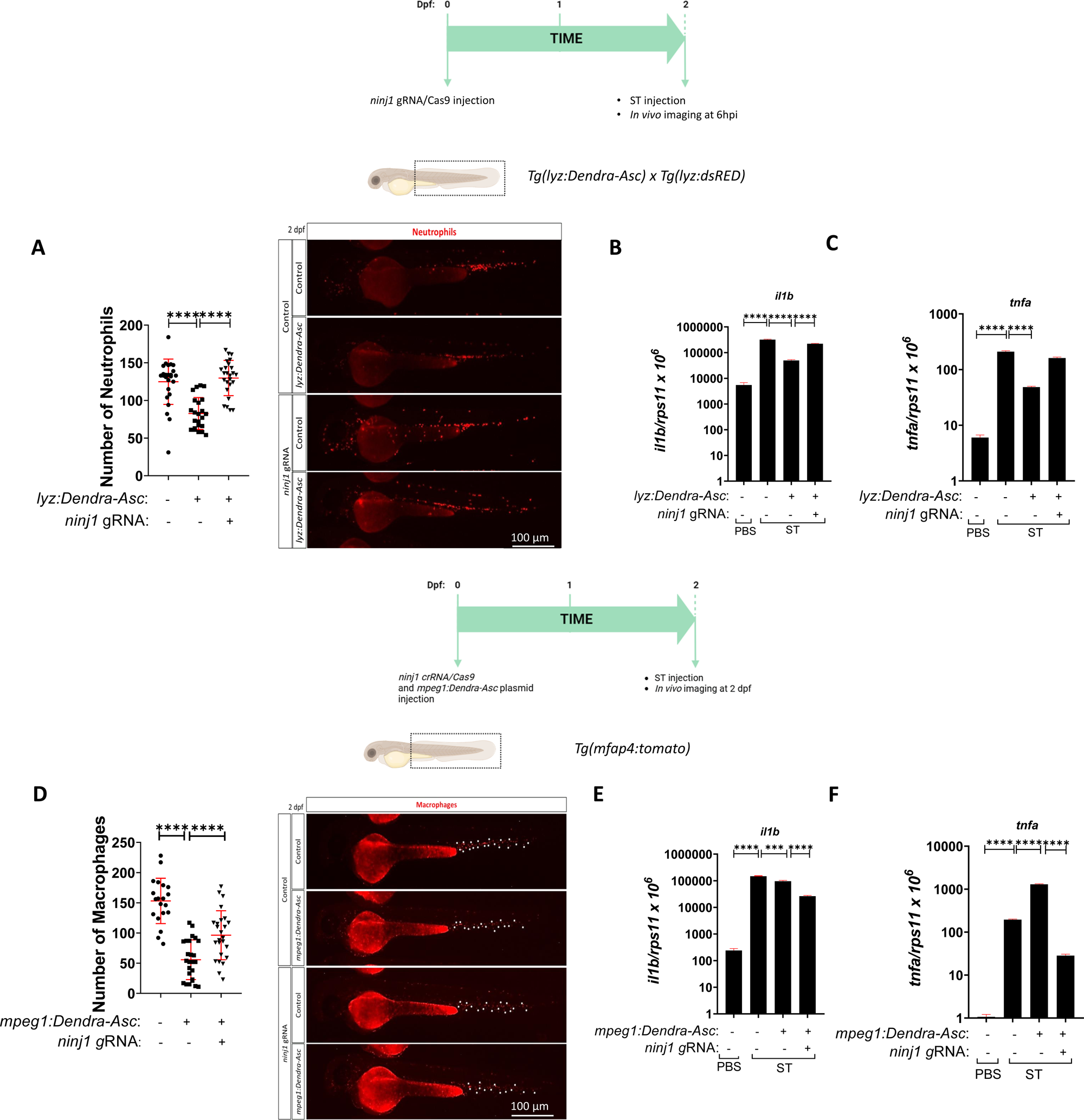
Ninj1-driven cell death mediate the crosstalk between neutrophils and macrophages. Zebrafish one-cell stage embryos were injected with *ninj1* gRNA/Cas9 complexes (**A-F**) or with *mpeg1:Dendra-Asc* plasmid (**D, E, F**). At 2 dpf larvae were injected with ST in the yolk sac (**B**, **C**, **E**, **F**). Neutrophils (**A**) and macrophages (**D**) in the body were visualized at 2 dpf by fluorescent microscopy. The transcript levels of the indicated genes (**B**, **C**, **E, F**) were analyzed at 24 hpi by RT-qPCR. Each dot represents one individual, and the means ± SEM for each group is also shown. *P* values were calculated using one-way analysis of variance (ANOVA) and Tukey multiple range test. \*\**P* ≤ 0.01, \*\*\**P* ≤ 0.001, and ****P ≤ 0.0001. The regions of interest used for quantification in all experiments are indicated in the schematic representation of the larvae. Bars: 100 µm.

**Figure S10 (related to Fig. 7).**
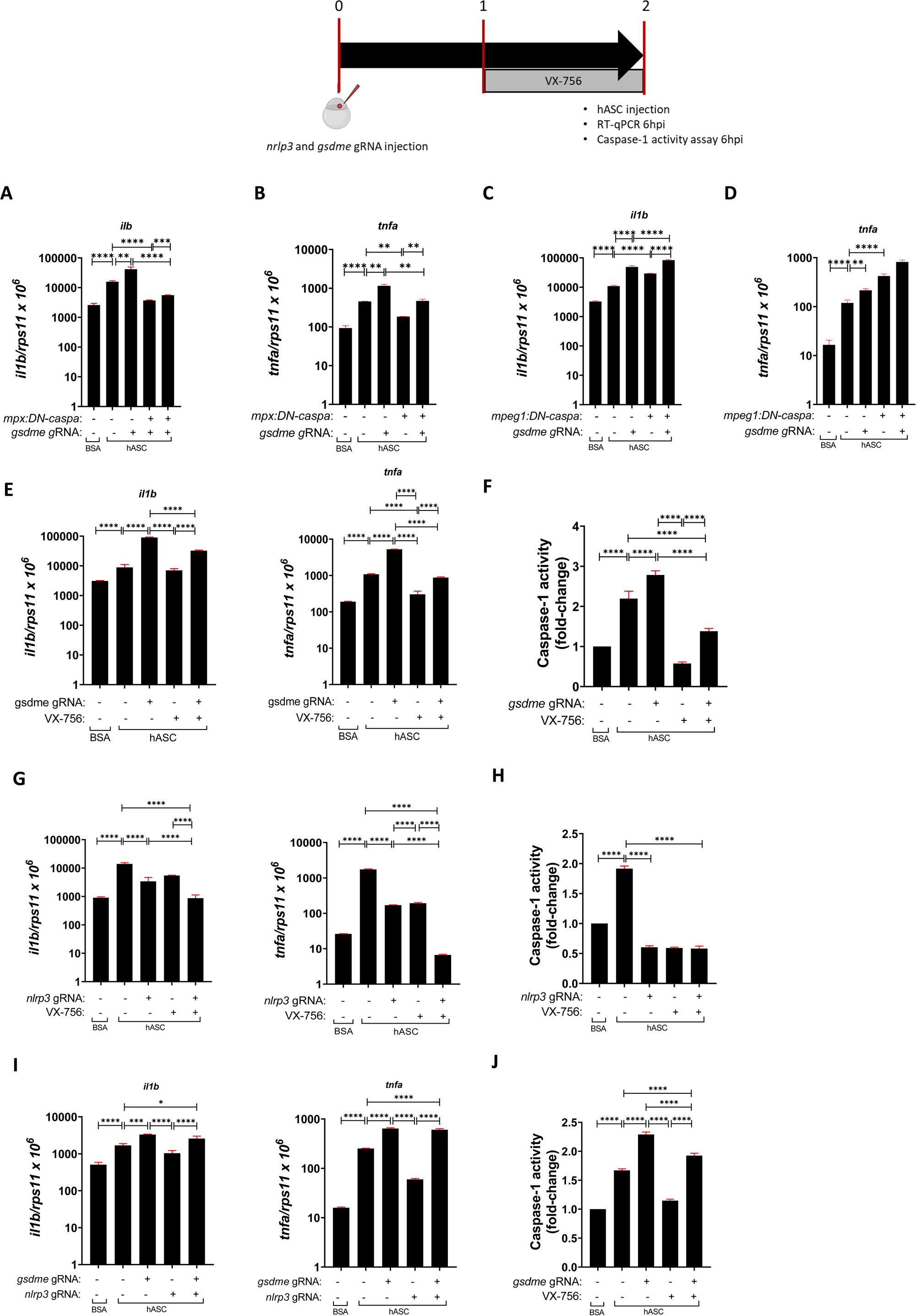
Gsdme drives the feedback inhibition of inflammasome activity in macrophages. Tg*(mpx:DN-caspa)* (**A, B**), Tg*(mpeg1:DN-caspa)* (**C, D**) embryos were used. Wild type (**E-J**) one-cell stage embryos were injected with *gsdme* and/or *nlrp3 g*RNAs/Cas9 complexes and treated with either DMSO or VX-765 from 1 to 2 dpf. At 1 dpf, either BSA or hASC were injected in the duct of Cuvier. The transcript levels of the indicated genes (**A-D, E, G, I**) were analyzed at 24 hpi by RT-qPCR. Caspase-1 activity was determined at 24 hpi using a fluorogenic substrate (**G, H**). Each dot represents one individual, and the means ± SEM for each group is also shown. *P* values were calculated using one-way analysis of variance (ANOVA) and Tukey multiple range test. \*\**P* ≤ 0.01, \*\*\**P* ≤ 0.001, and ****P ≤ 0.0001.

**Table S1.**
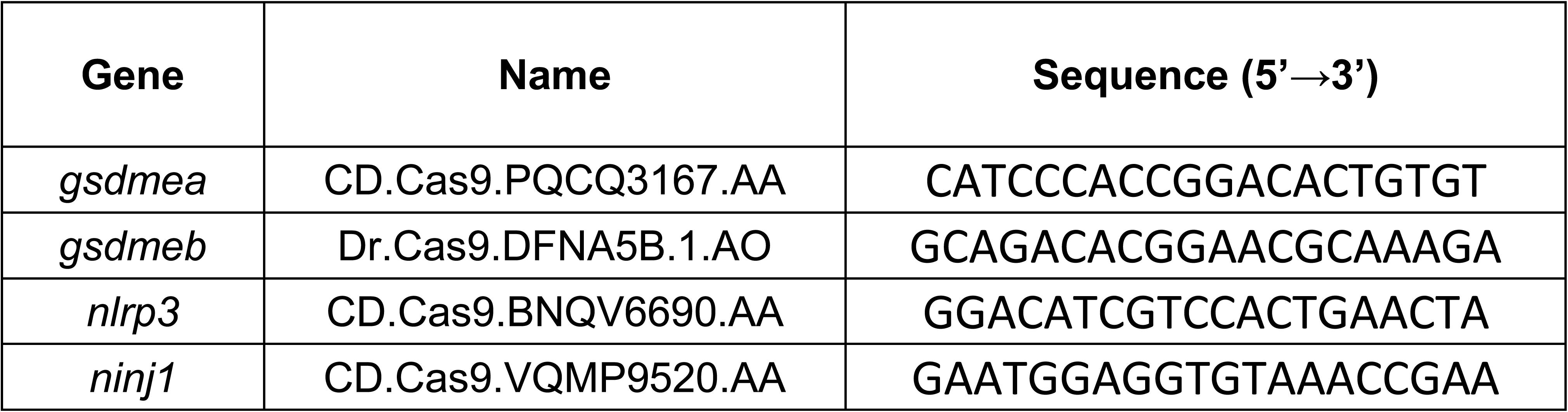
gRNA used in this study. The gene symbols followed the Zebrafish Nomenclature Guidelines (http://zfin.org/zf_info/nomen.html).

**Table S2.**
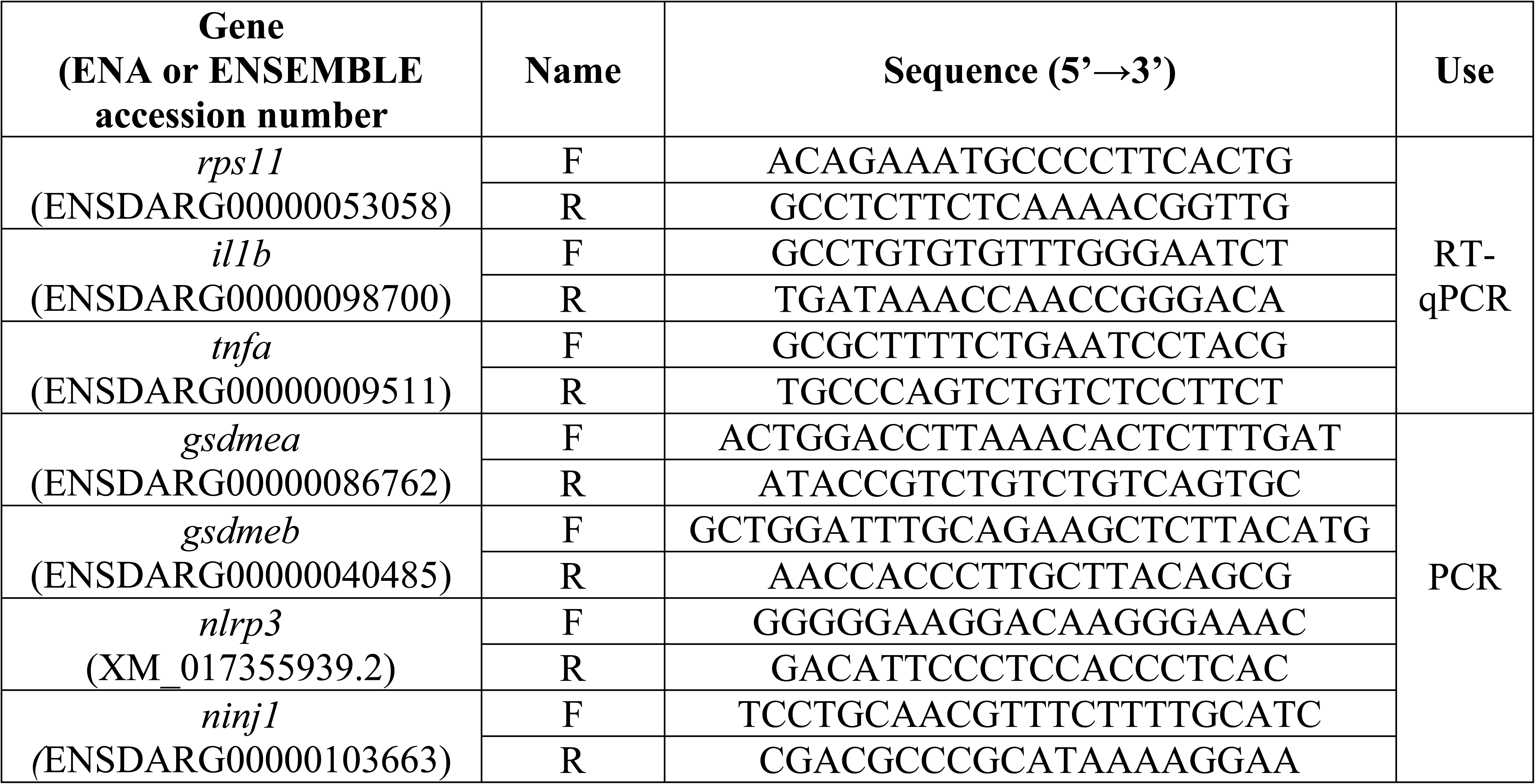
Primers used in this study. The gene symbols followed the Zebrafish Nomenclature Guidelines (http://zfin.org/zf_info/nomen.html).

## Video captions

**Videos 1 & 2 (related to Figure 6C): Neutrophil to neutrophil Asc speck transfer occurs *in vivo***. Vehicle (video 1) or recombinant S1WT (video 2) were injected in the hindbrain ventricle (HBV) of 2 dpf *Tg(lyz:Dendra-Asc)*. HBV photoconversion larvae was performed after injection using LSM880 airyscan and high resolution intravital imaging of CHT was performed for 8 hwith spinning disk confocal microscopy to visualize *in vivo* Asc-speck formation, release and spreading. Neutrophil are visualized in green and Asc-speck in red.

**Video 3 (related to Figure 6E): Neutrophil to macrophage Asc speck transfer occurs *in vivo***. Recombinant S1WT (+) or vehicle (−) were injected in the hindbrain ventricle (HBV) of 2 *Tg(lyz:Dendra-Asc)*; *Tg(mfap4:mCherry*). High resolution intravital imaging of CHT was performed with spinning disk confocal microscopy to visualize *in vivo* Asc-speck formation, release and spreading. Asc-speck were visualized in green and macrophages in red.

